# Simple study designs in ecology produce inaccurate estimates of biodiversity responses

**DOI:** 10.1101/612101

**Authors:** Alec P. Christie, Tatsuya Amano, Philip A. Martin, Gorm E. Shackelford, Benno I. Simmons, William J. Sutherland

## Abstract

1. Ecologists use a wide range of study designs to estimate the impact of interventions or threats but there are no quantitative comparisons of their accuracy. For example, while it is accepted that simpler designs, such as After (sampling sites post-impact without a control), Before-After (BA) and Control-Impact (CI), are less robust than Randomised Controlled Trials (RCT) and Before-After Control-Impact (BACI) designs, it is not known how much less accurate they are.
2. We simulate a step-change response of a population to an environmental impact using empirically-derived estimates of the major parameters. We use five ecological study designs to estimate the effect of this impact and evaluate each one by determining the percentage of simulations in which they accurately estimate the direction and magnitude of the environmental impact. We also simulate different numbers of replicates and assess several accuracy thresholds.
3. We demonstrate that BACI designs could be 1.1-1.5 times more accurate than RCTs, 2.9-4.1 times more accurate than BA, 3.8-5.6 times more accurate than CI, and 6.8-10.8 times more accurate than After designs, when estimating to within ±30% of the true effect (depending on the sample size). We also found that increasing sample size substantially increases the accuracy of BACI designs but only increases the precision of simpler designs around a biased estimate; only by using more robust designs can accuracy increase. Modestly increasing replication of both control and impact sites also increased the accuracy of BACI designs more than substantially increasing replicates in just one of these groups.
4. We argue that investment into using more robust designs in ecology, where possible, is extremely worthwhile given the inaccuracy of simpler designs, even when using large sample sizes. Based on our results we propose a weighting system that quantitatively ranks the accuracy of studies based on their study design and the number of replicates used. We hope these ‘accuracy weights’ enable researchers to better account for study design in evidence synthesis when assessing the reliability of a range of studies using a variety of designs.

## Introduction

Monitoring the impact of human activities on populations, communities and ecosystems is fundamental to understanding how to effectively protect and manage the natural world. This includes both monitoring the impacts of anthropogenic threats on biodiversity, as well as the effectiveness of management interventions to mitigate such threats. The main challenge with such monitoring is disentangling natural environmental change from anthropogenic change (Hewitt et al. 2001), whilst considering the focal impact’s statistical (Osenberg and Schmitt 1996, Stewart-Oaten 1996) and ecological significance (Wolfe et al. 1987). The complexity of natural ecosystems, including various sources of spatiotemporal variation and possible confounding variables, has led to considerable research effort to understand the best ways to study such impacts (Hewitt et al. 2001; Stewart-Oaten, Murdoch & Parker 1986; Osenberg et al. 2006; Block et al. 2001; McGarigal and Cushman 2002; Underwood & Chapman 2003). Improvements in study design have helped ecologists become better at accurately quantifying human impacts on biodiversity, yet a plethora of study designs with varying levels of complexity and associated biases persist (De Palma et al., 2018; Table 1).

**Table 1.**
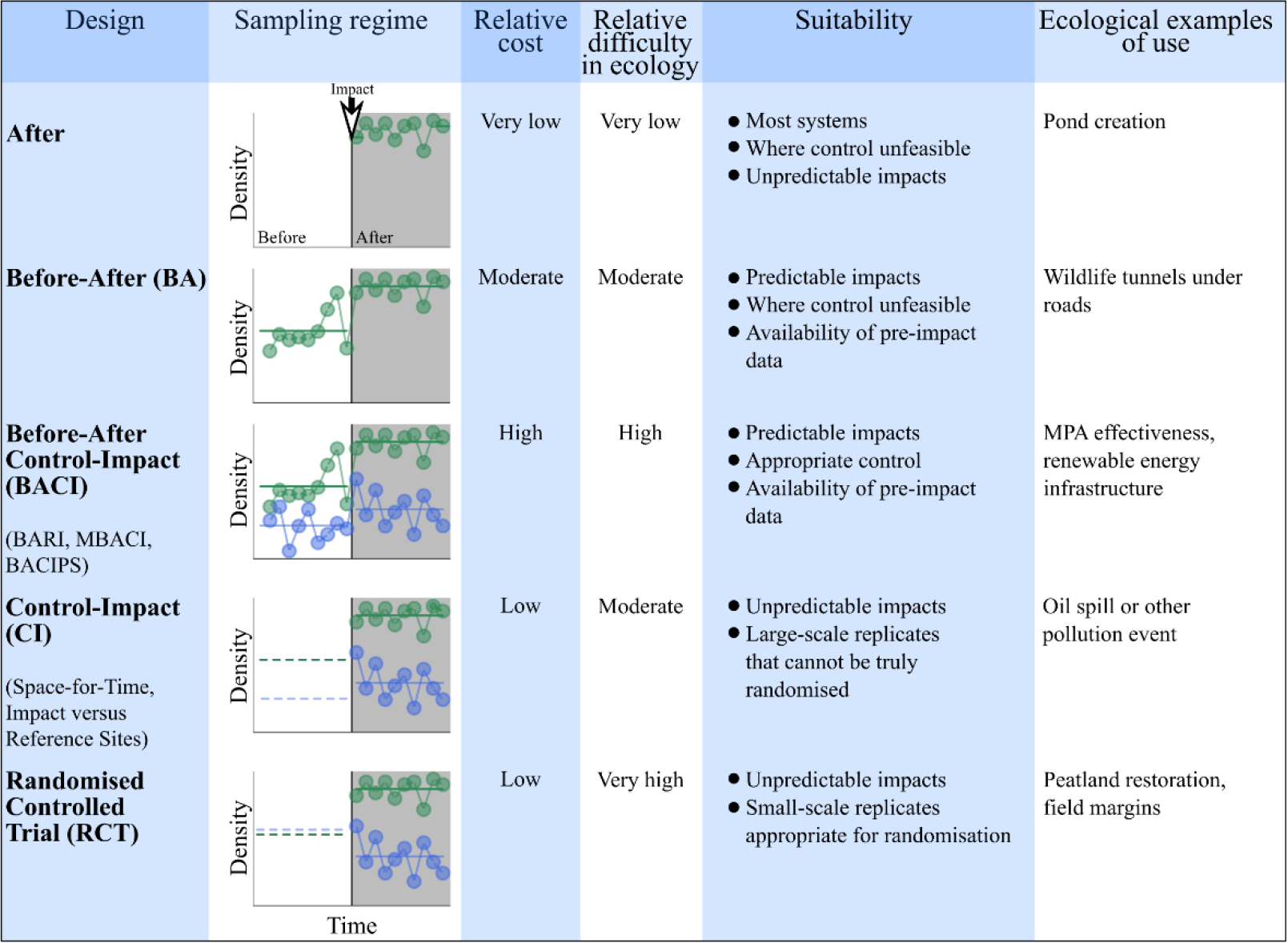
Comparison of the key features and characteristics of different study designs. Graphs show how designs sample impact (green points) and control (blue points) sites over time, before (white area) and after (grey area) an impact of a threat or intervention. Solid horizontal lines show the average density of the sites that is measured and used to calculate the effect size of each design. For CI and RCT designs, dashed horizontal lines show the average of control and impact sites before the impact - for RCTs these can be very similar (with high sample size), but for CI these can be very different. Note that many variants of the designs exist and the C for ‘Control’ can be replaced with R for ‘Reference (e.g. BARI or RI) depending on the circumstances (e.g. reference sites for restoration; Webb et al. 2012). The letter ‘M’ can also be used to indicate ‘Multiple’ sites for BACI/BARI designs (Downes et al. 2004).

| Design | Sampling regime | Relative cost | Relative difficulty in ecology | Suitability | Ecological examples of use |
| --- | --- | --- | --- | --- | --- |
| **After** |  | Very low | Very low | Most systems<br>Where control unfeasible<br>Unpredictable impacts | Pond creation |
| **Before-After (BA)** |  | Moderate | Moderate | Predictable impacts<br>Where control unfeasible<br>Availability of pre-impact data | Wildlife tunnels under roads |
| **Before-After Control-Impact (BACI)**<br>(BARI, MBACI, BACIPS) |  | High | High | Predictable impacts<br>Appropriate control<br>Availability of pre-impact data | MPA effectiveness, renewable energy infrastructure |
| **Control-Impact (CI)**<br>(Space-for-Time, Impact versus Reference Sites) |  | Low | Moderate | Unpredictable impacts<br>Large-scale replicates that cannot be truly randomised | Oil spill or other pollution event |
| **Randomised Controlled Trial (RCT)** |  | Low | Very high | Unpredictable impacts<br>Small-scale replicates appropriate for randomisation | Peatland restoration, field margins |

Study design is composed of three major aspects: (i) pre-impact sampling, (ii) use of controls, and (iii) randomised allocation of sample units (here we term these “sites”). Pre-impact sampling can be added to an After design - where a system is simply monitored after an impact of a threat or intervention occurs - to create a Before-After (BA) design (Table 1). This compares the system’s state before and after an impact, attempting to minimise bias from temporal variability and pre-impact conditions.

The addition of control sites to Before-After designs results in Before-After Control-Impact (BACI) designs, where the average difference between control and impact sites is compared before and after an intervention (Table 1; Stewart-Oaten, Murdoch & Parker 1986; Osenberg et al. 2006). BACI designs help to minimise bias from a lack of a control in simpler designs, using the difference in the before period between control and impact sites as a null hypothesis for the differences that would exist in the absence of the threat or intervention (Thiault et al. 2017). Problems associated with site-specific temporal variation in BACI studies can be addressed by sampling control and impact sites simultaneously, several times before and after the intervention (Before-After Control-Impact Paired-Series (BACIPS) design; Stewart-Oaten and Bence 2001).

Addition of randomly-allocated sites represents the final aspect of study design. Control-Impact (CI) designs, analogous with Space-For-Time Substitutions (França et al. 2016; De Palma et al. 2018) or Intervention Versus Reference Site designs (Stewart-Oaten and Bence 2001), compare control and impact sites after an impact has occurred. However, non-random site allocation can result in a violation of the assumption that the only differences between control and impact sites are as a result of the anthropogenic impact, leading to inaccurate results (De Palma et al. 2018; Damgaard 2019; Larsen et al. 2019; Table 1). Randomised Controlled Trials (RCTs) minimise this bias through randomised allocation of sites (Table 1) and, in theory, if sufficient sites and time steps are sampled to account for spatiotemporal variation, this reduces the need to sample before and after the intervention (i.e. using a BACI design) to account for any initial differences (Larsen et al. 2019; De Palma et al., 2018).

However, despite the generation of more robust approaches to quantifying impacts, greater usage of simpler, less robust designs persists. Three systematic maps on the biodiversity impacts of different threats and management interventions found that a low proportion of studies used BACI (6-29%) and BA designs (3-37%), but many more used CI designs (48-89%) (Bernes et al. 2015, Bernes et al. 2017, Papathanasopoulou et al. 2016).

The persistence of certain simpler designs probably reflects real-world constraints that prevent ecologists from using more robust study designs in the field. For example, RCTs are widely used in fields (e.g. medicine) where random allocation of small-scale experimental units to treatment groups is possible (Tugwell and Haynes 2006; Downs & Black 1998). However, RCTs often cannot be used in ecological studies (Stewart-Oaten & Bence 2001; Johnson 2002; Webb et al. 2012) because true randomisation of experimental units is much more difficult when using larger sites (e.g. protected areas) than when using smaller, more readily-available plots (Larsen et al. 2019). Therefore, ecologists tend to use pseudo-experimental designs that lack randomisation, such as BA, CI and BACI designs (Table 1; De Palma et al. 2018). However, cost and logistical constraints often prevent complex BACI and even simpler BA designs from being regularly used because of the need to revisit sites over several years (França et al. 2016, Osenberg et al. 2011; Table 1). Therefore, the greater prevalence of CI designs in the ecological literature probably reflects that they can be cheaper and easier to implement.

The disparities between the robustness of study designs and their usage is problematic because this may mean we are making misleading inferences about anthropogenic impacts. Although some empirical comparisons of the consequences of using BACI, BA and CI designs have been undertaken (Osenberg et al. 2011; França et al. 2016; Mahlum 2018), we still do not understand how much less accurate simpler designs are than more complex ones, or the influence on accuracy of increasing sample size of different study designs. A quantitative comparison of the accuracy of different study designs and their sample size would help us better understand these issues.

To address this knowledge gap, we simulate a hypothetical population’s response to an impact and compare how accurately different study designs estimate that impact’s effect. This novel simulation uses empirically-derived parameters (from 47 ecological datasets) to generate realistic control and impact data, before and after an impact. We use BACI, RCT, BA, CI and After designs with various levels of spatial replication (control and impact sites) to sample this simulated data and calculate an estimated effect size to compare against the true effect. We consider the accuracy of designs in terms of their ability to first, correctly predict the direction of the true effect and second, estimate the true effect to within a given percentage accuracy. We hope this study will enable the development of a quantitative hierarchy of the accuracy of different study designs, with major implications for future monitoring of anthropogenic impacts, as well as the use of ecological studies to inform policy and practice.

## Materials and methods

We simulated a system where the true density (*λ*) of a hypothetical population varied over a certain number of time steps (*T*) before and after a chronic impact occurred (Fig.1). For example, if *T*=10, then time steps 1 to 10 were classified as the ‘before period’ (i.e. before the impact occurred for control) and time steps 11 to 20 were classified as the ‘after period’ (Fig.1). The true density was monitored in sites where the impact occurred (‘impact sites’) and sites where the impact was absent (‘control sites’).

**Fig. 1.**
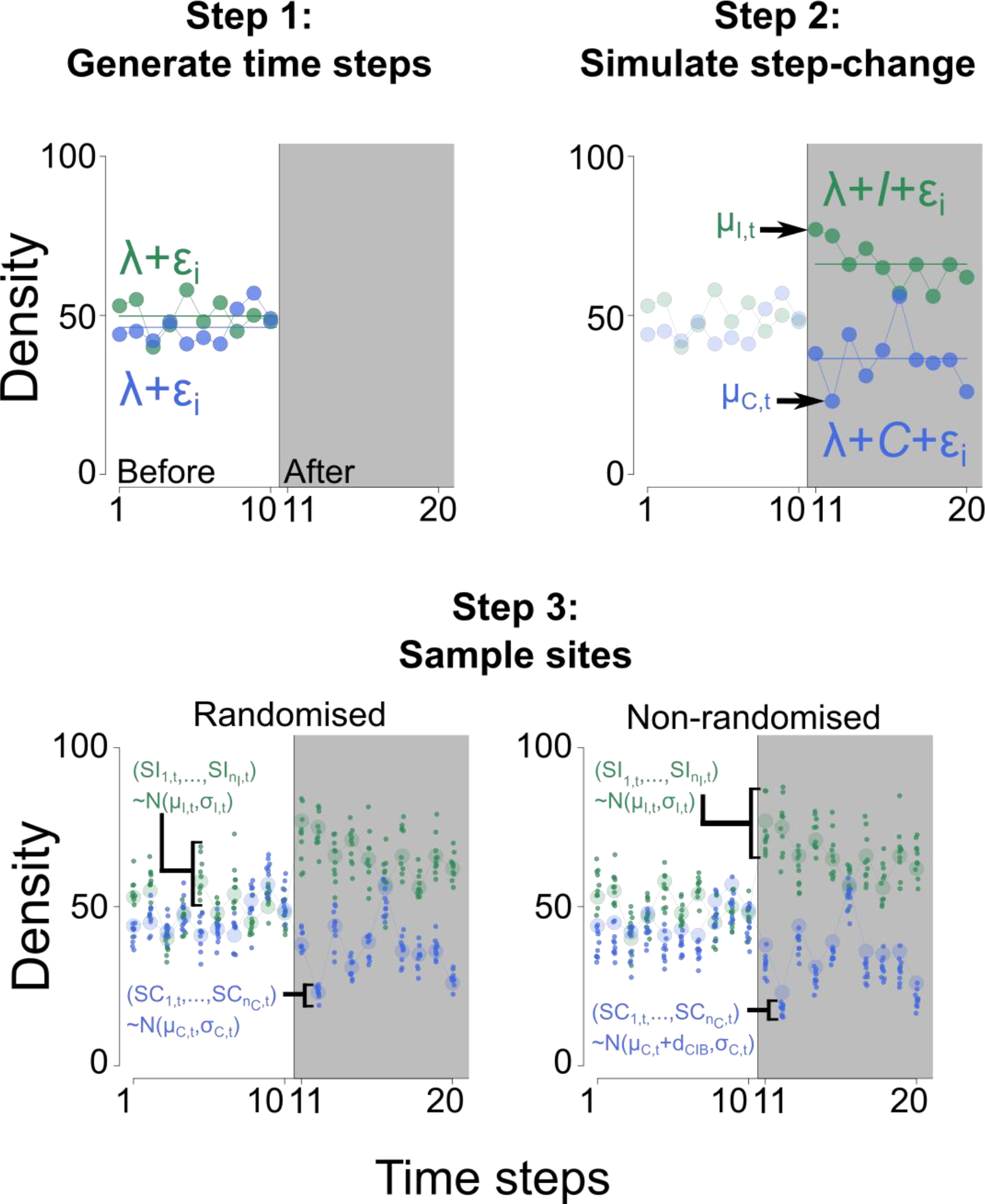
An overview of the simulation steps for both randomised and non-randomised design data. Step 1 shows true densities of control and impact sites generated in the before period (white area). Step 2 shows true densities of control and impact sites generated in the after period (grey area) to reflect a step-change response (using *I* and *C*). The true density in each time step (*t*) is shown (*μ*_*I*,*t*_, impact: green; and μ_*C*,*t*_, control: blue). Step 3 shows how control and impact sites (*SI* and *SC*) are sampled (*n*_*I*_ and *n*_*C*_ = 10) for both randomised and non-randomised designs.

We varied the true density (*λ*=50) over *T* time steps in the before period for control and impact sites by randomly sampling *T* values from a Poisson distribution. These *T* values defined the true density in each time step before the impact occurred (e.g. *μ*_*I*,*t*_for impact sites in the *t*th time step). To simulate a step-change response at both control and impact sites after the impact occurred (Fig.1), we sampled from a different Poisson distribution with *λ* adjusted by an empirically-derived amount *I* (*λ*+ *I*; Fig.1; Table 2) for impact sites and an empirically-derived amount *C* for control sites (*λ*+ *C*; Fig.1; Table 2). *I* and *C* were varied using empirical estimates of the proportional change in control and impact sites in the before period versus the after period, *p*_*I*_ and *p*_*C*_, respectively, sampled from 47 ecological datasets (*I*= *λ* · (*p*_*I*_ − 1); *C*= *λ* · (*p*_*C*_ − 1); Table 2; Supporting Information Section A).

**Table 2.** Definitions of all parameters used in the simulation and summary statistics for parameters that were empirically-derived (termed ‘emp.’; Supporting Information Section A). Equations show how each parameter was calculated. For empirically-calculated parameters, 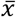 refers to the average of all sites of that type in that period (e.g. 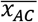 refers to the average of all control sites in the after period) taken from datasets (Supporting Information Section A). ‘sim.’ refers to parameters that are defined in the simulation.

| Parameter | Definition | Source | Equation | Mean | SD | Min | Max |
| --- | --- | --- | --- | --- | --- | --- | --- |
| $p_C$ | Change in control between before and after periods | Emp. | $p_C = \frac{\vert \bar{x}_{AC}\vert }{\vert \bar{x}_{BC}\vert }$ | 0.918 | 0.181 | 0.605 | 1.31 |
| $p_I$ | Change in impact between before and after periods | Emp. | $p_I = \frac{\vert \bar{x}_{AI}\vert }{\vert \bar{x}_{BI}\vert }$ | 0.967 | 0.230 | 0.579 | 1.46 |
| $p_{CIB}$ | Average value of control sites as a proportion of the average value of impact sites in the before period | Emp. | $p_{CIB} = \frac{\vert \bar{x}_{BC}\vert }{\vert \bar{x}_{BI}\vert }$ | 1.13 | 0.306 | 0.654 | 1.89 |
| $I$ | True change in impact sites from before to after impact | Emp. | $I = \lambda \cdot (p_I - 1)$ | -1.65 | 11.5 | -21.1 | 23.2 |
| $C$ | True change in control sites from before period to after period | Emp. | $C = \lambda \cdot (p_C - 1)$ | -4.10 | 9.05 | -19.8 | 15.4 |
| $d_{CIB}$ | Difference between true densities of control and impact sites in before period | Emp. | $d_{CIB} = \lambda \cdot (p_{CIB} - 1)$ | 6.60 | 15.3 | -17.3 | 44.5 |
| $\lambda$ | True density across all time steps | Fixed | $\lambda = 50$ | - | - | 50 | 50 |
| $T$ | Total number of time steps simulated | Sim. | $T = \{2,4,6,8,10\}$ | - | - | 2 | 10 |
| $n_T$ | Number of time steps sampled in each period | Fixed | $n_T = T$ | - | - | 2 | 10 |
| $\mu_{I,t}$ | True density in impact sites in time step $t$ | Sim. | Before: $\mu_{I,t} \sim \text{Poisson}(\lambda)$<br>After: $\mu_{I,t} \sim \text{Poisson}(\lambda + I)$ | - | - | - | - |
| $\sigma_{I,t}$ | Standard deviation of impact sites in time step $t$ | Sim. | $\sigma_{I,t} = CV_S \cdot \mu_{I,t}$ | - | - | - | - |
| $\mu_{C,t}$ | True density in control sites in time step $t$ | Sim. | Before: $\mu_{C,t} \sim \text{Poisson}(\lambda)$<br>After: $\mu_{C,t} \sim \text{Poisson}(\lambda + C)$ | - | - | - | - |
| $\sigma_{c,t}$ | Standard deviation of control sites in time step $t$ | Sim. | $\sigma_{c,t} = CV_S \cdot \mu_{c,t}$ | - | - | - | - |
| $CV_S$ | Coefficient of variation (variation amongst sites) | Sim. | $CV_S \sim N(\mu, \sigma, min, max)$ | 0.10 | 0.05 | 0 | 0.20 |
| $SI_{n,t}$ | $n^{th}$ impact site sampled in time step $t$ | Sim. | $(SI_{n,t}, \dots, SI_{n_I,t}) \sim N(\mu_{I,t}, \sigma_{I,t})$ | - | - | - | - |
| $SC_{n,t}$ | $n^{th}$ control site sampled in time step $t$ | Sim. | Randomised:<br>$(SC_{n,t}, \dots, SC_{n_C,t}) \sim N(\mu_{c,t}, \sigma_{c,t})$<br>Non-randomised:<br>$(SC_{n,t}, \dots, SC_{n_C,t}) \sim N(\mu_{c,t} + d_{CIB}, \sigma_{c,t})$ | - | - | - | - |
| $n_I$ | Number of impact sites sampled | Sim. | $n_I = \{1, 5, 10, 25, 50\}$ | - | - | 1 | 50 |
| $n_C$ | Number of control sites sampled | Sim. | $n_C = \{1, 5, 10, 25, 50\}$ | - | - | 1 | 50 |

While we concentrate on a step-change response in our simulation, temporal biodiversity dynamics following disturbances or interventions can follow different trajectories (Di Fonzo et al. 2013; Thiault et al. 2017). However, to simplify the simulation as much as possible, particularly in terms of computational demands, the use of a step-change response was most appropriate to test the relative accuracy of each design.

Using the simulated data for before and after periods we sampled various numbers of impact (*n*_*I*_) and control (*n*_*C*_) replicates (termed “sites” here but they could be plots or transects; Fig.1). For RCTs that use random allocation of sites to control and impact groups, we randomly sampled sites from two normal distributions for each time step: one with a mean, *μ*_*I*,*t*_, for impact sites and one with a mean, *μ*_*C*,*t*_, for control sites (Fig.1). The number of sites sampled was the same for all time steps. The standard deviation of each normal distribution represented the variation amongst sites and was calculated by multiplying the mean by the coefficient of variation (e.g. control sites: *σ*_*C*,*t*_= *μ*_*C*,*t*_ ⋅*CV*_*S*_; impact sites: *μ*_*I*,*t*_= *μI*,*t* ·*CV*; Table 2). We varied *CV*_*S*_ by randomly drawing values from a truncated normal distribution: N(*μ* = 0.1, *σ* = 0.05, *min* = 0, *max* = 0.2).

To account for non-random allocation of sites to control and impact groups in the other designs (BACI, BA, CI, After), we repeated the same approach but with one important modification. We adjusted the true density of control sites in every time step, *μ*_*C*,*t*_, by an empirically-derived amount, *d*_*CIB*_ (*μ*_*C*,*t*_+ *d*_*CIB*_; Fig.1; Table 2). To vary *d*_*CIB*_, we used empirical estimates of the proportional difference between control and impact sites in the before period, *p*_*CIB*_, sampled from 47 ecological datasets (*d*_*CIB*_= λ ·(*p*_*CIB*_-1); Table 2; Supporting Information Section A). This simulated difference between control and impact sites accounted for different levels of site selection bias in non-randomised designs, including the situation where little or no bias may be present (e.g. *d*_*CIB*_≈0).

We calculated effect sizes for each design by first finding the mean density of sampled sites across all time steps for control and impact groups in the before period (*Before*_*Impact*_, *Before*_*Control*_) and the after period (*After*_*Impact*_, *After*_*Control*_). We assumed that sampling occurred in all time steps (*n*_T_ = *T*) in the before and after periods. We did this as the investigator may wish to only estimate the effect over a certain timescale (which will be context-specific) and we lacked the computational capabilities to simulate all possible sampling permutations using fewer than the full number of time steps (e.g. deciding which time steps to sample in certain intervals; e.g. Wauchope et al. 2019).

Effect sizes were calculated using these mean densities, as appropriate for each study design (Table 3). For example, the effect size for an RCT was found by subtracting *After*_*Control*_ from *After*_*Impact*_, whilst the effect size for a BA design was found by subtracting *Before*_Impact_ from *After*_*Impact*_ (Table 3). The exception was the After design, for which we found the mean of sampled sites in the first time step of the after period and subtracted this from the mean of sampled sites in the final time step (Table 3). We defined the true effect as the change in the true density of impact sites between the before and after periods minus the equivalent change in the true density of control sites (Table 3). As discussed previously, we did this because we wanted to compare each design’s relative accuracy at estimating the true effect over the number of time steps we simulated.

**Table 3.**
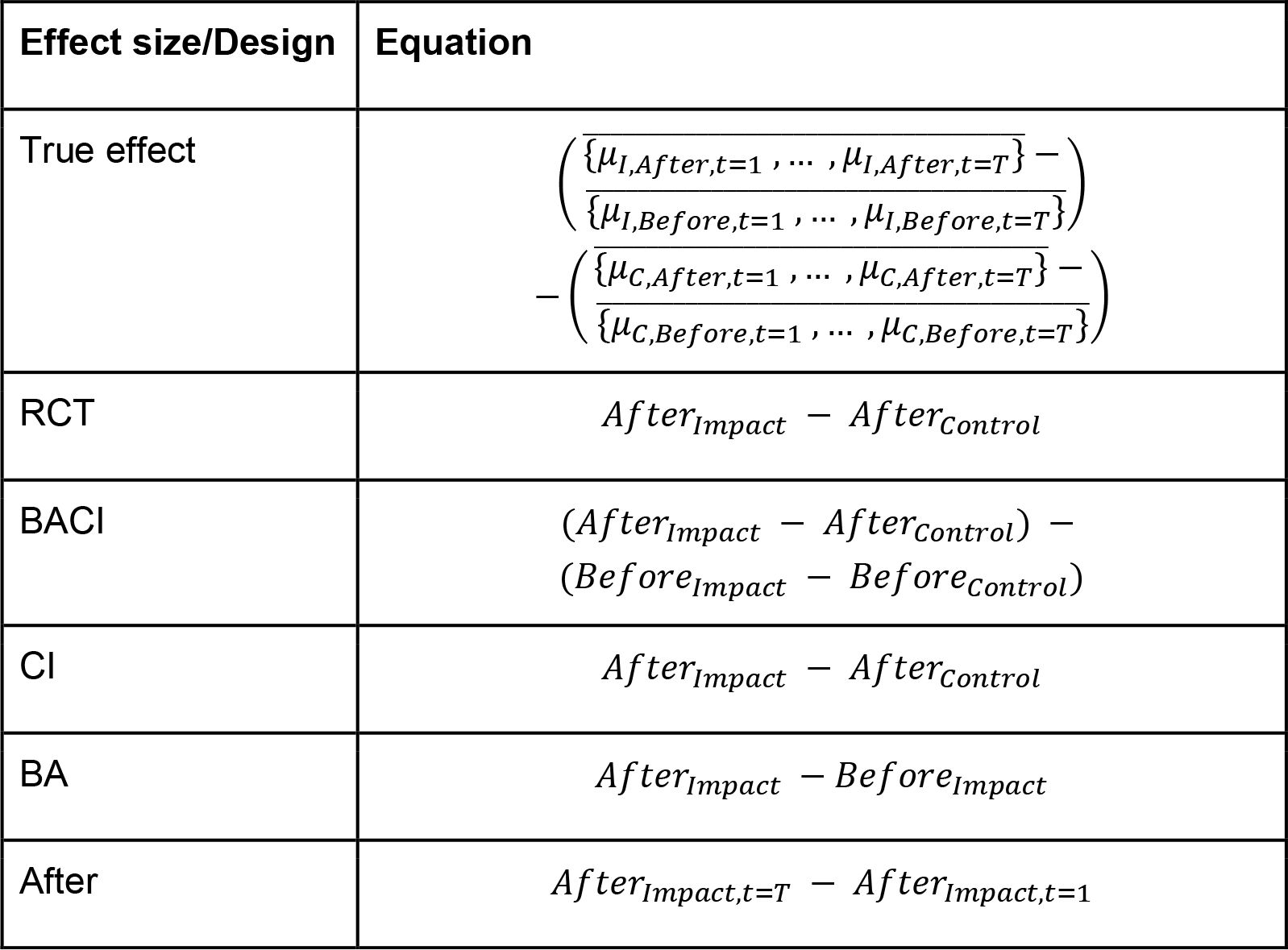
Equations for how effect sizes were calculated for different designs using mean densities of control or impact sites in each period (e.g. *After*_*Impact*_ refers to the mean of sampled impact sites across all time steps in the after period). For the After design, the effect size was calculated by finding the difference between the final time step (t=*T*) and the first time step of the after period (1).

We ran the simulation under 1000 different scenarios, varying: (i) the true change in control sites (*C*); (ii) the true change in impact sites (*I*); (iii) the mean difference between control and impact sites in the before period (*p*_*CIB*_); and (iv) the variation between sites (*CV*_*S*_). For each simulation scenario, we varied the number of time steps simulated (*T* = 2, 4, 6, 8 or 10), as well as the number of impact sites (*n*_*I*_ = 1, 5, 10, 25, 50) and control sites (*n*_*C*_ = 1, 5, 10, 25, 50) sampled independently to use every possible pairwise combination - this resulted in a total of 125 combinations. Overall, we simulated 1000 scenarios with 125 different sampling combinations in each, repeating each scenario 1000 times (1000 × (1000 × 125) = 1.25 × 10^8^ runs).

The effect sizes for each design from the simulation were used to investigate the relative accuracy of the different designs. To do this we calculated the percentage of simulation repetitions in which each design’s effect size: (i) had the same direction as the true effect size; and (ii) was both within a given percentage of true effect size and of the same direction. We believe these two metrics capture the major aspects of accuracy that are desirable in a study design.

We calculated both percentages for all possible pairwise combinations of control and impact sites (e.g. one control and one impact site, one control and five impact sites etc.) and the second percentage for five accuracy thresholds (within ±10, 20, 30, 40 and 50% of true effect size). We used Generalised Linear Models with a binomial error family to determine the relationship between the accuracy of each design (the response variable; see below) and the three explanatory variables (number of control sites, number of impact sites and the accuracy threshold). We did not include control sites as an explanatory variable for BA and After designs.

For the response variable we used the number of simulation runs that were successes and the number of simulation runs that were failures; a success was defined as when the estimated effect size was within a given percentage of the true effect size and of the same direction. Based on graphical observations of the relationship between the response and explanatory variables (Fig.3), we log transformed each of these explanatory variables sequentially, creating several models (Supporting Tables 1-5). We calculated McFadden’s pseudo-R squared (McFadden 1974) for each model and picked the best model using the lowest AIC value (Supporting Tables 1-5).

We used R statistical software version 3.5.1 (R Core Team 2018) with the doParallel package (Microsoft Corporation & Weston 2017) to increase computational performance. R code to perform the simulations and analyses is provided in Supporting Information.

## Results

There was large variation in the performance of study designs in correctly estimating the direction of the effect. As overall patterns were similar across simulations with different time steps (see Supporting Figures 1 & 2), here we present results when six time steps were simulated in both the before and after periods. BACI designs were best at correctly identifying the direction of the true effect size (in at least 92.9% of simulation repetitions; Fig.2), followed by RCTs (at least 91.5% of the time). Both BACI and RCTs far outperformed CI, BA, and particularly After designs; CI designs performed slightly better than BA designs (approximately 71.7% versus 68.3%) and both performed far better than After designs, which were worse than random chance with an accuracy of approximately 49.9% (Fig.2).

RCTs, After, BA and CI designs showed negligible improvement in correctly identifying the direction of the effect with increasing number of impact and/or control sites (increases from one control and one impact site to 50 control and 50 impact sites: After = +0.0%; BA = +0.6%; CI= +0.1%; RCT= +1.3%; Fig.2). However, there were small asymptotic increases in performance with increasing sample size for BACI designs (+5.2% from one control and impact site to 50 control and impact sites; Fig.2).

**Fig. 2.**
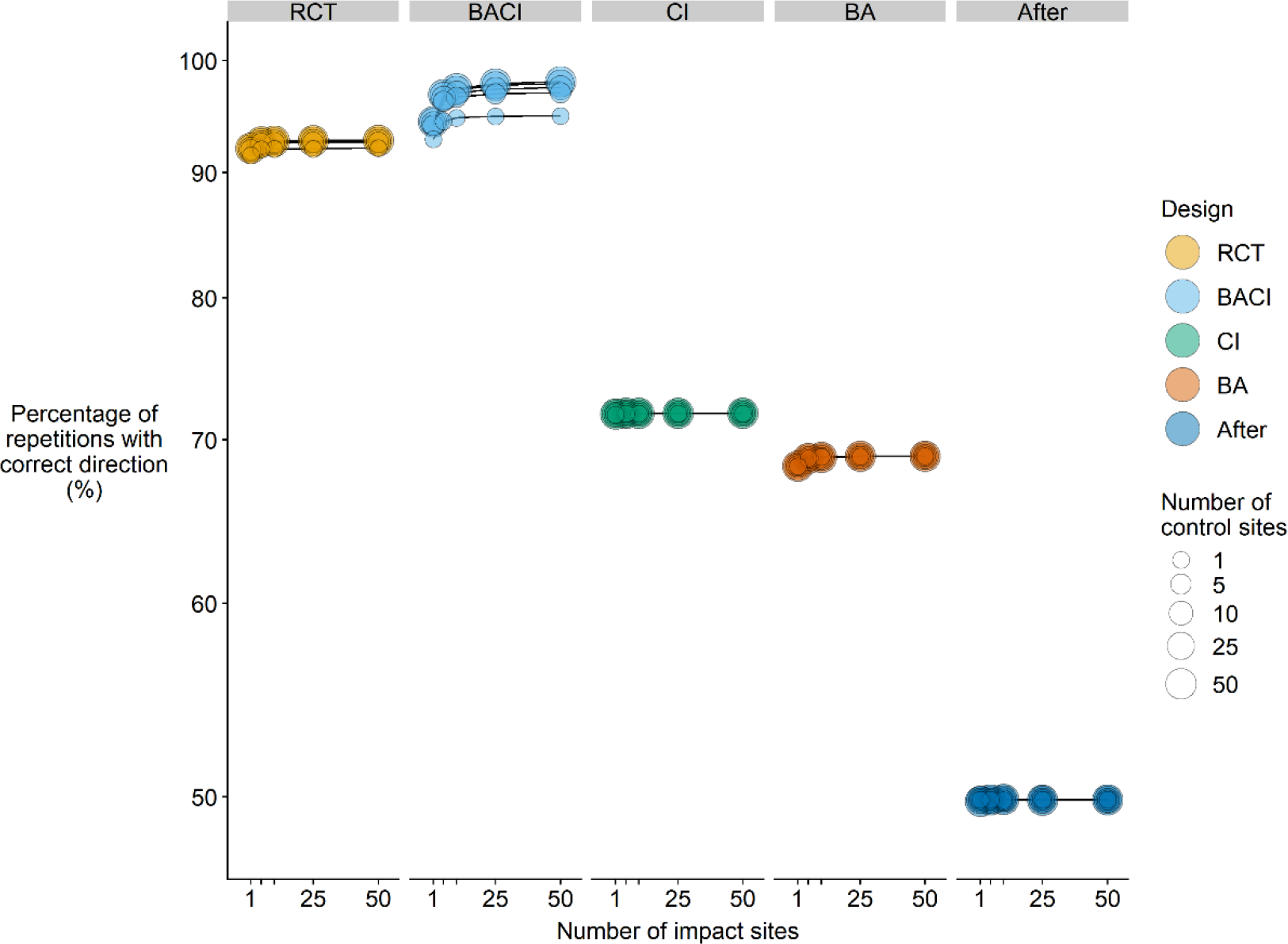
Accuracy of different designs measured by the percentage of simulation repetitions in which the design’s effect size had the correct direction for multiple levels of spatial replication (control and impact sites separately). Y-axis uses a log_10_ scale and circle size denotes the number of control sites. See Table 1 for the definition of each design.

We see similar patterns when considering the percentage of repetitions for which the effect size was both within a certain percentage of the true effect size and had the correct direction (Fig.3). First, RCT and BACI designs are still clearly far more accurate at estimating the true effect size than CI, BA or After designs (for 30% accuracy threshold: BACI ≥ 60.9%, RCT ≥ 53.1%, BA ≥ 20.8%, CI ≥ 15.9%, After ≥ 8.3%; Fig.3). Second, changing the percentage threshold is highly influential and as the threshold becomes greater (within a greater % of the true effect size) accuracy increases substantially for all designs (Fig.3; see also Supporting Figure 2). Third, BACI accuracy increased to a much greater extent with increasing spatial replication than for other designs which showed limited improvements (Fig.3). For the 30% accuracy threshold, increasing replication from one control and impact site to 50 control and impact sites resulted in an increase of 28.8% for BACI compared to +5.7% for RCTs, +0.8% for BA, +0.2% for CI and −0.7% for After (Fig.3).

**Fig. 3.**
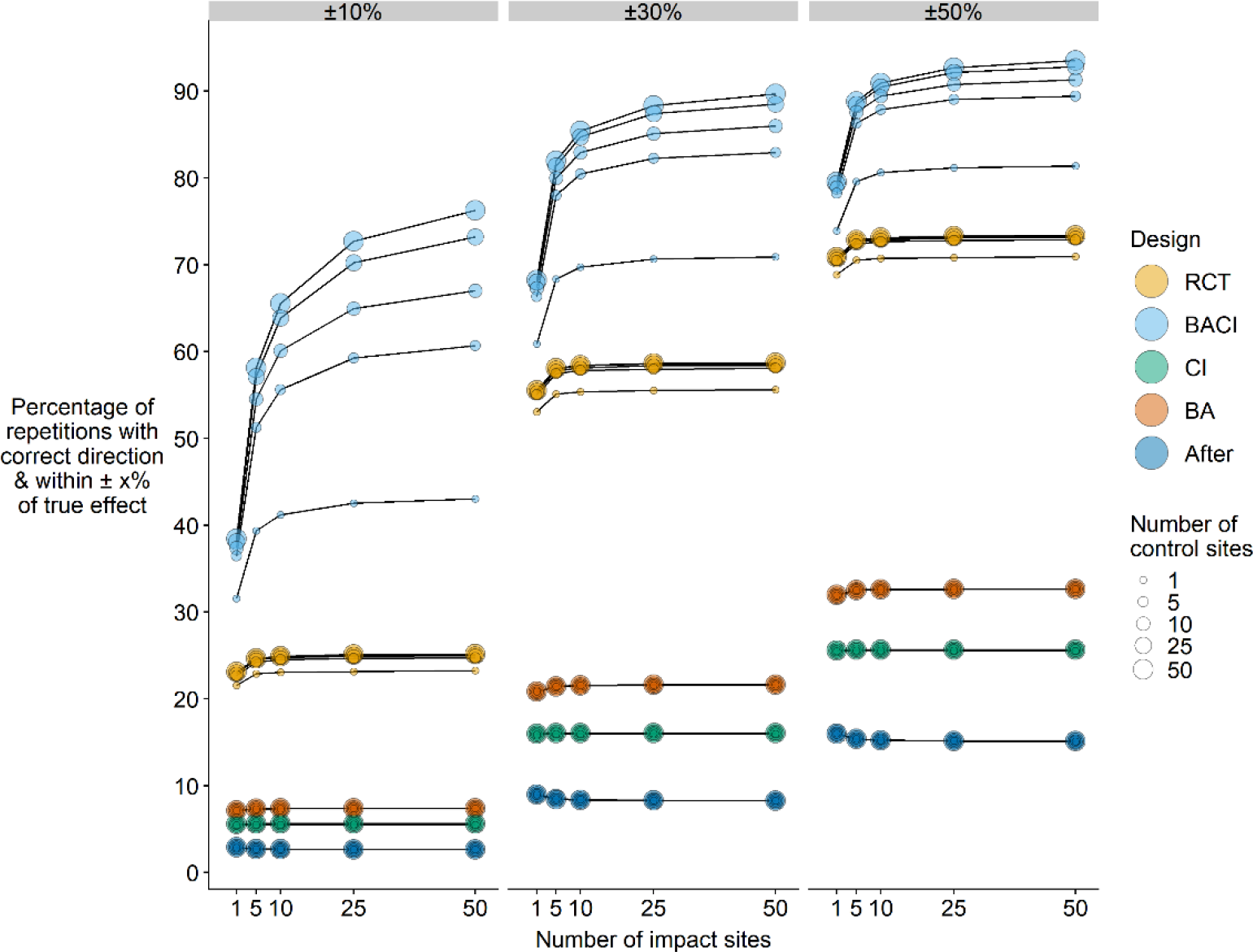
Accuracy of different designs measured by the percentage of simulation repetitions in which the design’s effect size was both within 10, 30 or 50% of the true effect size and had the correct direction for multiple levels of spatial replication (control and impact sites separately). Circle size denotes the number of control sites. See Table 1 for the definition of each design.

Moreover, increasing replication moderately in both control and impact sites resulted in greater accuracy than only increasing replication in either impact or control sites for BACI designs (Fig.3). For example, using a 30% accuracy threshold, the accuracy of BACI designs using five impact and five control sites was greater than using 50 impact sites and one control site (78.0% versus 70.9%).

While BA designs were poorer at correctly predicting the direction of the effect than CI designs, they appeared to be more accurate when we accounted for the proximity of their effect sizes to the truth. Furthermore, as the accuracy threshold increases from 10% to 50%, BA designs shifted from being almost as accurate as CI designs (~2% more) to being moderately more accurate (~7% more; Fig.3). Similarly, both BA and CI designs became relatively more accurate compared to After designs with an increasing accuracy threshold (Fig.3).

Using Generalised Linear Models to determine factors affecting the accuracy of each design, the full model (i.e. the model with all explanatory variables considered) showed the smallest AIC in all designs (Table 4; Supplementary Tables 1-5). The estimated coefficients for the number of control and impact sites were smaller for After, BA, CI and RCT designs compared to BACI designs, suggesting its greater importance in BACI designs. The level of accuracy (A) was an important predictor of the accuracy of all designs (Table 4). For CI, BACI and RCT designs, there was little difference in the importance of control versus impact sites in predicting accuracy. Very high Pseudo-R^2^ values showed that our models explained far greater levels of variation in the data than null models (Table 4).

**Table 4.** Results of Generalised Linear Models determining factors that affect the accuracy of each study design based on simulation results - coefficients are in log odds (3.s.f.). McFadden’s pseudo-R squared (McFadden 1974) was calculated. *n*_*I*_ = number of independent impact units, *n*_*C*_ = number of independent control units, A = accuracy threshold as a proportion (e.g. 0.3 = ±30%).

| | Intercept | | Accuracy threshold<br>$\ln(A)$ | | Number of impact sites<br>$\ln(n_I)$ | | Number of control sites<br>$\ln(n_C)$ | | |
| --- | --- | --- | --- | --- | --- | --- | --- | --- | --- |
| Design | Coef. | SE | Coef. | SE | Coef. | SE | Coef. | SE | Pseudo-R<sup>2</sup> |
| *RCT* | 1.73 | 6.67<br>$\times 10^{-5}$ | 1.31 | 3.56<br>$\times 10^{-5}$ | 0.0297 | 1.41<br>$\times 10^{-5}$ | 0.0287 | 1.41<br>$\times 10^{-5}$ | 0.999 |
| *BACI* | 1.54 | 7.15<br>$\times 10^{-5}$ | 1.07 | 3.74<br>$\times 10^{-5}$ | 0.272 | 1.59<br>$\times 10^{-5}$ | 0.226 | 1.59<br>$\times 10^{-5}$ | 0.951 |
| *CI* | -0.323 | 8.75<br>$\times 10^{-5}$ | 1.10 | 5.39<br>$\times 10^{-5}$ | 1.84<br>$\times 10^{-3}$ | 1.85<br>$\times 10^{-5}$ | 1.89<br>$\times 10^{-3}$ | 1.85<br>$\times 10^{-5}$ | 0.999 |
| *BA* | 0.0277 | 7.02<br>$\times 10^{-5}$ | 1.13 | 4.80<br>$\times 10^{-5}$ | 0.0104 | 1.68<br>$\times 10^{-5}$ | - | - | 1.000 |
| *After* | -0.868 | 9.98<br>$\times 10^{-5}$ | 1.19 | 7.31<br>$\times 10^{-5}$ | -0.0203 | 2.36<br>$\times 10^{-5}$ | - | - | 0.997 |

Converting estimated coefficients (β) from log odds (Table 4) to probabilities gives an ‘accuracy weight’ for each design (see discussion for how these weights could be applied):

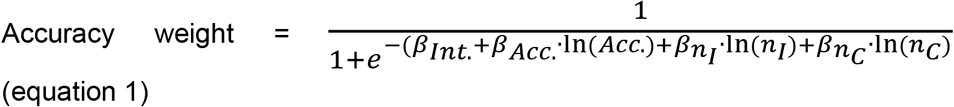

where *β*_*Int*._ = Intercept coefficient, *β*_*ACC*._ = Accuracy threshold coefficient, *β*_*n*_*I*__= Impact sites coefficient and *β*_*n*_*C*__= Control sites coefficient.

The accuracy weight can be calculated for any study using information about its study design and the number of control and impact units used. If a study design does not use Control sites (n_C_), then this part of the equation is simply omitted. For example, a study by Merz & Chan (2005) which used a BA design with seven impact units would receive an accuracy weight of:

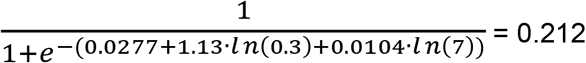

(assuming an accuracy threshold of ±30% (0.3); for estimated coefficients see BA, Table 4).

A more robust study by França et al. (2016) used a BACI design, 29 impact and five control units and thus receives an accuracy ‘weight’ of:

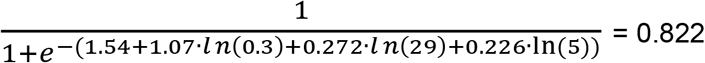

(see BACI, Table 4).

Similarly, the accuracy weight for a study by Westera et al. (2003) which used a CI design with three control and three impact units equals:

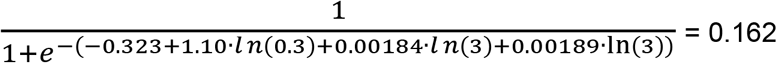

(see CI, Table 4).

## Discussion

Using this simulation we have demonstrated that BACI and RCT designs are far more accurate than BA, CI and After designs. We showed that BACI and RCT designs, depending on the level of spatial replication, identified the correct direction of the change ~20-30% more often than CI and BA designs, and ~40-45% more than After designs. At an accuracy threshold of ±30%, BACI designs were also 1.1-1.5 times more accurate than RCTs, 2.9-4.1 times more accurate than BA, 3.8-5.6 times more accurate than CI, and 6.8-10.8 times more accurate than After designs (depending on the sample size). This is because, although it is easy to increase precision in After, BA and CI designs by simply increasing sample size, these designs usually yield biased estimates. Greater accuracy is best achieved by using more robust designs that remove biases, such as BACI and RCTs, with greater numbers of replicates. BACI designs appeared to be more accurate than RCTs, probably because RCTs can only minimise initial differences between control and impact sites, but unlike BACI designs cannot completely account for them.

Our results provide strong evidence that simpler study designs (e.g. BA and CI) often yield different effect sizes and inferences to BACI designs, as observed empirically by previous studies (Osenberg et al. 2011; França et al. 2016; Mahlum 2018). Therefore, studies using After, BA and CI designs seriously risk presenting misleading conclusions on the impact of threats and interventions. To our knowledge, this simulation is not only the first quantitative comparison to demonstrate this, but also to show *how* much less accurate BA and CI designs may be on average when using ecologically-realistic parameters under varying levels of spatial replication.

Our quantitative results provide a general overview of the relative accuracy of different study designs in ecology but will exhibit context-dependency for different fields of ecology. However, as our simulation can be parameterised using empirically-derived estimates, if sufficient empirical data can be collected from different fields of ecology, future research could use our R code to examine the context-dependency of our results.

Another key finding with major implications is that increasing sample size generally has no impact on the accuracy of simpler study designs. This is because unless the assumptions of these designs hold true (e.g. no change in the control for BA designs, or the only differences between control and impact sites are as a result of the anthropogenic impact for CI designs), increasing sample size will only increase precision around a biased estimate of the true effect of the impact. We have confidence in this conclusion given that we used empirically-derived parameter values from 47 ecological datasets to effectively estimate the likelihood and magnitude of any violation of these assumptions in ecological studies. Therefore, our results provide strong evidence that, generally in ecology, investing sampling effort into replicating simpler designs is inefficient and would be better spent on implementing more robust designs whenever possible.

However, we realise that these more complex designs will not always be easy to implement. Nevertheless, we argue that there are still situations where ecologists can use these designs and yet fail to do so; promoting greater awareness of more robust designs and opportunities where they can be used is important. For example, where prior knowledge exists of when the impact of an intervention or threat may occur (or where suitable data pre-impact data is available retrospectively), ecologists should try to use BACI designs (e.g. infrastructure projects, Protected Area designation). If this is not feasible, but small plot-based experimental units can be truly randomised to treatments and control (e.g. field margins, grassland plots), RCTs could be used. We also acknowledge that the expensive nature of BACI designs, due to the need for multiple visits to study sites before and after the impact, does hamper their implementation and requires action from the scientific community (De Palma et al. 2018). However, we must consider the trade-off between using more expensive, robust study designs and the costs of using potentially misleading studies with simpler, biased designs as evidence to inform important policy and practice. Therefore, we argue that since long-term monitoring is often incompatible with short-term timescales of existing funding models (Osenberg et al. 2011; De Palma et al. 2018), longer-term funding to facilitate the use of more robust designs is needed.

Unfortunately, it seems the use of simpler designs is likely to persist for the foreseeable future, so we argue that our results also have major implications for evidence synthesis. We argue that our quantitative comparison of the accuracy of study designs could help to develop novel methods and weighting schemes for synthesising studies that vary markedly in their design and quality (Spake and Doncaster 2017). Conventional meta-analysis typically tries to account for the quality of primary studies, using inverse variance as weights (Marín-Martínez and Sánchez-Meca 2010, Koricheva and Gurevitch 2014). However, this could greatly reduce the number of primary studies that can be included, as not all studies report variance (Koricheva and Gurevitch 2014, Stewart 2010). Alternative approaches of evidence synthesis to tackle poor data reporting, such as non-parametric weighting by sample size, have been encouraged (Gurevitch and Hedges 1999) and proposed (Mayerhofer et al. 2013, Adams et al. 1997) but these weights fail to consider wider aspects of study quality, including pseudoreplication and study design (Spake and Doncaster 2017). Whilst recent efforts for assessing evidence quantitatively by study design (Norris et al. 2012, Webb et al. 2012, Mupepele et al. 2016, Mupepele & Dormann 2017) are welcomed, their weights are relatively simplistic and discrete (e.g. simple integer scores or categories) and do not seem to have been informed using quantitative evidence.

The relationships we have presented between accuracy, study design and sample size can be used to generate a continuous weighting scale, accounting for many more aspects of study quality than are considered by solely weighting studies by sample size or variance. Using our results (Table 4) we can produce ‘accuracy weights’ (eqn 1), which could be used to modulate study weights in meta-analysis - i.e. giving greater influence to studies with more accurate designs (see examples in Results).

Although study design and spatial replication only assess part of study quality (Spake and Doncaster 2017), our weights are also versatile and could be modulated using subject-specific quality checklists to help incorporate more context-specific factors of study quality into evidence assessment such as size of sampling unit, temporal replication and internal validity (Spake and Doncaster 2017; Mupepele et al. 2016; Bilotta et al. 2014). For example, our weights could be multiplied by the percentage of criteria met in quality checklists. However, adding extra components to the evidence assessment process also needs to be balanced against the effort expended in doing so.

More generally, our weights could also be used to assign studies into different accuracy categories, giving a rapid, easily interpretable way to communicate the robustness of evidence to decision-makers - e.g. in evidence toolkits such as Conservation Evidence (2019) or Higgins et al. (2014). For example, each study could be assigned a weight and then the average weight could be found for a set of studies to indicate the relative strength of the evidence (e.g. mean weight ≥0.8 = strong evidence). Systematic reviews could be weighted using this method, either by totalling the weights of studies they review or by finding the median or mean weight of these studies. We welcome future research to explore how best to apply and incorporate these weights into evidence assessment.

Future research could also seek to explore other aspects of study design, such as considering different types and intensities of temporal replication (e.g. time intervals between sampling; Wauchope et al. 2019; or pseudoreplication in time; Stewart-Oaten, Murdoch and Parker 1986). The relative performance of designs under different types of responses to anthropogenic impacts (e.g. sigmoidal, linear or asymptotic; Thiault et al. 2017) or lag periods (De Palma et al. 2018) could also be investigated further. Our simulation code could be easily modified to facilitate the exploration of these important issues (Supporting Information).

Overall, we have quantitatively shown for the first time how much less accurate simpler study designs are compared to more complex ones, generating a far better quantitative understanding of the relative accuracy of different study designs. Our accuracy weights could also offer a powerful, yet versatile new approach to weighing up evidence from studies with different designs, with major implications for the future of evidence synthesis. We hope our work encourages greater discussion of study design in ecology and demonstrates that we need to tackle the serious consequences of using different study designs to make inferences in ecology.

## Acknowledgements

For providing empirical data used in parameterising simulation, we thank: Anna A. Sher, Ricardo Rocha, Annelies de Backer, Aurora Torres, Carlos Palacín, Juan Carlos Alonso, Barry Baldigo, Brendan P Kelaher, Daniel Mateos, Doriane Stagnol, Dominique Davoult, Filipe Machado França, Heather Major, Ian Jones, Jake Bicknell, Jenyffer Vierheller Vieira, Maria Carmen Ruiz-Delgado, Ruben Heleno, Joachim Claudet, Kade Mills, Kevin Stokesbury, Bradley Harris, Mehdi Adjeroud, Michael Craig, Michele Meroni, Norbertas Noreika, Janne S Kotiaho, Patrick Edwards, Rafael Barrientos, Carlos Ponce, Carlos A Martín, Beatriz Martín, Ricardo Ceia, Roland Pitcher, Sarah Clarke, Oliver Tully, Shailesh Sharma, Just Cebrian, Thomas Stanley, Tyler Eddy, Jonathan Gardner, Anjali Pande, Adrià López-Baucells, Christoph Meyer, Alvaro Antón, Bob McConnaughey, Corrine Watts, David Abecasis, Luciana Cibils, Monica Montefalcone, Teppo Vehanen, Aki Mäki-Petäys, Ari Huusko, Juan Jacobo Schmitter-Soto, Matt Rinella, Garth Hodgson, Hartwell Welsh, Mikael van Deurs, Mary Donovan, Axel Schwerk, Jill Shaffer, Deborah A Buhl, Alberto Velando, Dolores River Restoration Partnership, Javier Seoane Pinilla, Andrew Page, Matt Dasey, David Maguire, Jos Barlow, Júlio Louzada, Rachel T Buxton, Carley R Schacter, Melinda G Conners, Koniambo Nickel, Ginger Soproner, CSIRO, Arturo Elosegi, Loreto García-Arberas, Joserra Díez, Ana Rallo. Datasets found freely available online: Burge et al. (2017), Dietl & Durham (2016), Sepúlveda & Valdivia (2016), Williams et al. (2014), Moland et al. (2013). Thanks also to Grania Smith for helping to proofread manuscripts. Author funding sources: TA was supported by the Grantham Foundation for the Protection of the Environment, the Kenneth Miller Trust and the Australian Research Council Future Fellowship (FT180100354); WJS, PAM and GES are supported by Arcadia and The David and Claudia Harding Foundation; BIS and APC were supported by the Natural Environment Research Council as part of the Cambridge Earth System Science NERC DTP [NE/L002507/1].

## Author contributions

APC, WJS and TA conceived the ideas and designed simulation methodology; APC analysed the data and led the writing of the manuscript. All authors contributed critically to the development of methodology and manuscript drafts, as well as giving final approval for publication.

## Data Accessibility

R script and empirical data used in simulations uploaded as Supporting Information.

## Supporting Information

Supporting information contains two files: (1) a PDF containing supporting figures and explanations of empirically-derived parameters; and (2) a zip file containing simulation code to reproduce methods and empirical data used to parameterise simulation.

