## Supporting Code for "Simple study designs in ecology produce inaccurate estimates of biodiversity responses": reproducecode.R

```
##### R libraries required
library(truncnorm)
library(doParallel) ####if you can run in parallel (i.e. your machine has enough cores)

#### First read in empirical dataset which has parameter values

#cht = proportional change in impact
#chc = proportional change in control
#tcd = proportional difference between impact and control (impact-control) in before period

paramdis = read.csv(choose.files(),stringsAsFactors = FALSE)
head(paramdis)
str(paramdis)

paramdis = data.frame(paramdis)
paramdis$Chc = as.numeric(paramdis$Chc)
paramdis$Cht = as.numeric(paramdis$Cht)
paramdis$tcd= as.numeric(paramdis$tcd )

runs=1000 ### number of unique runs in simulation
multb1 = 1000 #### number of repetitions of each run

### find 25% and 75% quantiles for each parameter
quantschc = quantile(paramdis$Chc,na.rm=TRUE)[c(2,4)]
quantscht =quantile(paramdis$Cht,na.rm=TRUE)[c(2,4)]
quantstcd =quantile(paramdis$tcd,na.rm=TRUE)[c(2,4)]

### restrict parameter values to InterQuartile Range
Chc2 = paramdis$Chc[which(paramdis$Chc>=quantschc[1] & paramdis$Chc<=quantschc[2])] 
Cht2 = paramdis$Cht[which(paramdis$Cht>=quantscht[1] & paramdis$Cht<=quantscht[2])]
tcd2 = paramdis$tcd[which(paramdis$tcd>=quantstcd[1] & paramdis$tcd<=quantstcd[2])] 

#### sample from parameter values 1000 times with replacement to generate 1000 unique parameter runs
#### then repeat these 1000 times (1000 repetitions)
cht = rep(round(sample(Cht2[which(is.na(Cht2)==FALSE)],runs,replace=TRUE),digits=3),multb1)
chc = rep(round(sample(Chc2[which(is.na(Chc2)==FALSE)],runs,replace=TRUE),digits=3),multb1)
tcd = rep(round(sample(tcd2[which(is.na(tcd2)==FALSE)],runs,replace=TRUE),digits=3),multb1)

### sample from truncated normal distribution to get values of between site coefficient of variation (CV) for each of the 1000 runs

svar1= rep(rtruncnorm(runs,0,0.2,0.1,0.05),multb1)

### put parameters togethere in dataframe ready for simulation
params = data.frame(cbind(chc,cht,tcd,svar1))


#### function for simulation

###zeta = repetition number

supersimsc <- function(zeta){
  
  #### input parameters from dataframe we created
  svar=params$svar1[zeta]
  
  ### set lambda to 50 for population abundance
  startvalt = 50
  
  cht1 = (params$cht[zeta]-1)*startvalt
  chc1 = (params$chc[zeta]-1)*startvalt
  tcd1 = (params$tcd[zeta]-1)*startvalt
  
  
  ### Possible combinations of sites and time steps
  sitesvals1t = c(1,5,10,25,50) ###possble numbers of impact sites sampled
  sitesvals1c = c(1,5,10,25,50) ###possble numbers of control sites sampled
  yrst = c(2,4,6,8,10) ###possble numbers of time steps sampled and simulated
  ##### beware amount of memory this will require, data produced = ~9GB!
  
  ### create a matrix to capture effect sizes for each design and all possible combinations of spatial replication (5x5) and time steps sampled (x4)
  results = matrix(ncol=9,nrow=length(sitesvals1t)*length(sitesvals1c)*length(c(2,4,6,8,10)))
  z=1
  
  for (tt1 in 1:length(yrst)){ ###loop through all possible numbers of time steps simulated
    
  ###set number of time steps to simulate in each period 
  ###(i.e. if maxyears = 10, 10 before + 10 after = 20 time steps total)
  maxyears = yrst[tt1]
  
  ### first generate time steps for impact and control group in before period
  ### then generate same number of time steps, but with a lambda adjusted by cht1 and chc1 (change in impact and control) 
  year1tat = c(rpois(maxyears,startvalt),rpois(maxyears,startvalt+cht1))
  year1tac = c(rpois(maxyears,startvalt),rpois(maxyears,startvalt+chc1))
  
  ### loop through all these combinations of sites and time steps sampled and generate effect sizes
  
  for(st in 1:length(sitesvals1t)){
    for(sc in 1:length(sitesvals1c)){
      
        s1=sitesvals1t[st] # no. impact sites sampled
        s2=sitesvals1c[sc] # no. control sites sampled
        tt=yrst[tt1] # no. time steps sampled
        
        #### create matrices to capture site data for each time step and group
        
        ### randomised controlled trial design 
        temptranci = matrix(ncol=tt,nrow=s1)### After Impact
        tempcranci = matrix(ncol=tt,nrow=s2) ### After Control
        
        ### non-randomised designs: baci (using all 4 matrices) or BA (using only first 2 matrices)
        ### or CI (using 2nd and 4th matrices) or After (using only 2nd matrix)
        temptb = matrix(ncol=tt,nrow=s1) ### Before Impact
        tempta = matrix(ncol=tt,nrow=s1) ### After Impact
        tempcb = matrix(ncol=tt,nrow=s2) ### Before Control
        tempca = matrix(ncol=tt,nrow=s2) ### After Control
        
        for(t1 in 1:tt){ ### for every time step, sample sites
          
          #### use pmax to prevent negative values
          #### generate sites using normal distribution with a mean taken from the time step value
          #### svar is the coefficient of variation (CV) we parameterised earlier and is multiplied by the mean value to give the sd of the normal distribution
          
          ### randomised controlled trial design 
          temptranci[,t1] = pmax((rnorm(s1, year1tat[maxyears+t1],year1tat[maxyears+t1]*svar)),0) #After impact
          tempcranci[,t1] = pmax((rnorm(s2, year1tac[maxyears+t1],year1tac[maxyears+t1]*svar)),0) #After control
          
          ### After or CI or Before-After or BACI design data
          ###note addition of tcd1 for control data to reflect non-random site selection so 
          ###impact and control groups differ on average by tcd1
          
          tempta[,t1] = pmax((rnorm(s1, year1tat[maxyears+t1],year1tat[maxyears+t1]*svar)),0) #After impact
          temptb[,t1] = pmax((rnorm(s1, year1tat[maxyears-t1+1],year1tat[maxyears-t1+1]*svar)),0) #Before impact
          
          tempca[,t1] = pmax((rnorm(s2, pmax(year1tac[maxyears+t1]+tcd1,0),pmax(year1tac[maxyears+t1]+tcd1,0)*svar)),0) #After control
          tempcb[,t1] = pmax((rnorm(s2, pmax(year1tac[maxyears-t1+1]+tcd1,0),pmax(year1tac[maxyears-t1+1]+tcd1,0)*svar)),0) #Before control
          
        }
        
        ### find means for impact and control in each period and use to calculate effect size for each design
        ### store them along with how many time steps and sites were sampled and repeat
        
        results[z,1] <- mean(temptranci) - mean(tempcranci) ###RCT
        results[z,2] <- mean(tempta) - mean(temptb)  ### BA
        results[z,3] <- mean(tempta) - mean(tempca) ### CI
        results[z,4] <- (mean(tempta) -  mean(tempca)) - (mean(temptb)-mean(tempcb))  ###BACI
        results[z,5] <- (mean(tempta[,tt]) -  mean(tempta[,1])) ### After using final time step sampled - first time step sampled
        results[z,6] <- tt ###number of time steps sampled
        results[z,7] <- s1 ###number of impact sites sampled
        results[z,8] <- s2 ###number of control sites sampled
        
        ###true change over simulated number of time steps
        results[z,9] <- (mean(year1tat[(maxyears+1):(maxyears+tt)]) -  mean(year1tat[(maxyears-tt+1):maxyears])) -
                        (mean(year1tac[(maxyears+1):(maxyears+tt)])-mean(year1tac[(maxyears-tt+1):maxyears]))
                        
        
        z=z+1
      }
    }
  }

  ###package up results
  return(list(cbind(results)))
  
}

### test how long it will take serially (without parallelisation) - test with 100 repetitions and multiply up to get a rough estimate of total time (multiply time by 10000)
samp = sample(1:(nrow(params)),100,replace=FALSE)

system.time({
  sapply(samp,supersimsc)
})


### we ran this in parallel on a windows machine with the following code
ncores = 8 ### put in the number of cores you want to use - on our Hyper-threaded Core i9 (20 core) PC 
           ###we found the best performance was achieved at 8 cores (probably as true number of cores is 10)

c1 <- makeCluster(ncores) ###create a cluster of cores
clusterExport(c1,c('params')) ### import the empirical dataset that it will need to run simulation
clusterEvalQ(c1,library('truncnorm')) ### import any libraries it may need

y <- (1:nrow(params)) ###find number of repetitions (1x10^6)

system.time(
  devall <- parSapplyLB(c1,y,supersimsc) #### time it with system.time if desired and run in parallel with load balancing (LB)
)

stopCluster(c1) ### make sure to stop cluster at the end

### parallel processing generates list output so we need to rbind this together to get results
devall2 = do.call(rbind, devall)

str(devall2) #check it worked

### label columns
colnames(devall2) = c("RCT",
                      "BA","CI","BACI","After",
                      "yr1","tsi1","csi1","Truth")
###write output to a csv
#write.csv(devall2,"simresults.csv",row.names=FALSE)


#############################################################
###### Analysis #############################################
#############################################################

library(readr)
library(data.table)
library(cowplot)
library(ggplot2)
library(viridis)

## if you want to reimport csv file of results
#devall2= fread(choose.files())

str(devall2)
devall3=data.frame(devall2)
str(devall3)

### indexes for each combination of number of sites and yrs
sivalst = c(1,5,10,25,50)
sivalsc = c(1,5,10,25,50)
yrvals = c(2,4,6,8,10)

##### Function to generate percentage of repetitions in which effect size of a design was the same sign (+,- or 0) as the true effect size

percor = 1
percorfunc = function(zeta) {
  percor = (length(which(devall2[,zeta] > 0 & devall2[,9] >0)) + length(which(devall2[,zeta] == 0 & devall2[,9] ==0)) + length(which(devall2[,zeta] < 0 & devall2[,9] <0)))/
    length(devall2[,zeta])
  return(percor)}


#### loop through all number of site combinations and find percentage with correct sign
z=1
res=matrix(ncol=8,nrow=length(sivalst)*length(sivalsc)*length(yrvals))
for(t in 1:length(yrvals)){
  for(i in 1:length(sivalst)){
    for(p in 1:length(sivalsc)){
      devall2 = devall3[which(devall3$tsi1== sivalst[i]&devall3$csi1== sivalsc[p] &devall3$yr1==yrvals[t]),]
      
      res[z,1:5] = sapply(1:5,percorfunc)*100
      res[z,6] = sivalst[i]
      res[z,7] = sivalsc[p]
      res[z,8] = yrvals[t]
      z=z+1
    }
  }
}

des1=list()
for(i in 1:5){
  des1[[i]] = rep(i,length(sivalst)*length(sivalsc)*length(yrvals))
}
des1=unlist(des1)

###reorder data to plot and join with which design is which
res3all3 = data.frame(cbind(c(res[,c(1,4,3,2,5)]),rep(res[,6],5),rep(res[,7],5),rep(res[,8],5),des1))
head(res3all3)
colnames(res3all3) = c("dev","sit","sic","yr","des1")
res3all3$sit=as.numeric(res3all3$sit)
res3all3$sic=as.factor(res3all3$sic)
res3all3$des1=as.factor(res3all3$des1)
levels(res3all3$des1)=c("RCT","BACI","CI","BA","After")

###reorder by the number of control sites sampled
res3all3 = res3all3[rev(order(res3all3$sic)),]

### colour-blind palette to use (Link: https://www.nature.com/articles/nmeth.1618)
clrs=c(rgb(230/255,159/255,0),rgb(86/255,180/255,233/255),rgb(0,158/255,115/255),rgb(213/255,94/255,0),rgb(0,114/255,178/255))

### subset for number of time steps = 6
res3all4 = subset(res3all3,res3all3$yr==6)

### plot line and point graph with point size as proportional to number of control sites sampled
ggplot(res3all4,aes(x=sit,y=dev,group=interaction(des1,sic))) + geom_line()+ 
  geom_point(aes(size=sic,fill=des1),pch=21,alpha=0.5)  + 
  facet_wrap(des1~.,nrow=1)+
  scale_x_continuous(name="Number of impact sites",limits=c(-5,55),breaks=c(1,5,10,25,50),labels=c(1,"","",25,50))+
  scale_y_continuous(name="Percentage of\nrepetitions with\ncorrect direction\n(%)",breaks=seq(50,100,10),labels=seq(50,100,10),limits=c(48,100))+
  coord_trans(y = "log10")+ theme(axis.title.y  = element_text(angle=0, vjust=0.5, size=15),strip.text.x = element_text(size = 15),axis.text.x=element_text(size=15),axis.text.y=element_text(size=15),axis.title.x=element_text(size=15,hjust=0.5),legend.text = element_text(size=15),legend.title=element_text(size=15))+
  theme(aspect.ratio=5,panel.background=element_blank(),panel.grid.major=element_blank(),panel.grid.minor=element_blank(),plot.margin = unit( c(0.1,0.1,0.1,0.1) , "in" ) )+
  scale_colour_manual(name="Design",values=clrs) +scale_fill_manual(name="Design",values=clrs) +guides(fill = guide_legend(override.aes = list(size=10)))+
  scale_size_manual(name="Number of\ncontrol sites",values=5:9)

### save your graph as a png
#ggsave(filename = "results1.png",dpi = 300,width = 30,height = 30,units = "cm")

###suppl.info graph
ggplot(res3all3,aes(x=sit,y=dev,group=interaction(des1,sic))) + geom_line()+ 
  geom_point(aes(size=sic,fill=des1),pch=21,alpha=0.5)  + 
  facet_wrap(yr~.,nrow=1)+
  scale_x_continuous(name="Number of impact sites",limits=c(-5,55),breaks=c(1,5,10,25,50),labels=c(1,"","",25,50))+
  scale_y_continuous(name="Percentage of\nrepetitions with\ncorrect direction\n(%)",breaks=seq(50,100,10),labels=seq(50,100,10),limits=c(48,100))+
  coord_trans(y = "log10")+ theme(axis.title.y  = element_text(angle=0, vjust=0.5, size=15),strip.text.x = element_text(size = 15),axis.text.x=element_text(size=15),axis.text.y=element_text(size=15),axis.title.x=element_text(size=15,hjust=0.5),legend.text = element_text(size=15),legend.title=element_text(size=15))+
  theme(aspect.ratio=3,panel.background=element_blank(),panel.grid.major=element_blank(),panel.grid.minor=element_blank(),plot.margin = unit( c(0.1,0.1,0.1,0.1) , "in" ) )+
  scale_colour_manual(name="Design",values=clrs) +scale_fill_manual(name="Design",values=clrs) +guides(fill = guide_legend(override.aes = list(size=10)))+
  scale_size_manual(name="Number of\ncontrol sites",values=5:9)

#ggsave(filename = "supplinfographsdirection.png",dpi = 300,width = 30,height = 30,units = "cm")


##### find the number of simulation repetitions where both sign was correct and the effect size of a design was within a given % of the true effect size

z1=1
res3all=list()

for(a in 1:5){ ### a determines accuracy threshold (e.g. 10% to 50% in steps of 10%)
  percor = 1
  percorfunc = function(zeta) {
    percor = (length(which((abs(devall2[,zeta]) >= abs(devall2[,9]*(1-(a/10))) & abs(devall2[,zeta]) <= abs(devall2[,9]*(1+(a/10)))) & ((devall2[,zeta] > 0 & devall2[,9] >0) | (devall2[,zeta] == 0 & devall2[,9] ==0) | (devall2[,zeta] < 0 & devall2[,9] <0)))))/
      length(devall2[,zeta])
    return(percor)}
  
  res2=matrix(ncol=8,nrow=length(sivalst)*length(sivalsc)*length(yrvals))
  z=1
  for(t in 1:length(yrvals)){
    for(i in 1:length(sivalst)){
      for(p in 1:length(sivalsc)){
        devall2 = devall3[which(devall3$tsi1== sivalst[i]&devall3$csi1== sivalsc[p] & devall3$yr1==yrvals[t]),]
        
        res2[z,1:5] = sapply(1:5,percorfunc)*100
        res2[z,6] = sivalst[i]
        res2[z,7] = sivalsc[p]
        res2[z,8] = yrvals[t]
        z=z+1
      }
    }
  }
  
  res3all[[z1]] = cbind(res2,a/10)
  z1=z1+1
}

### recombine results from list

res3all2 = do.call(rbind, res3all)

des1=list()
for(i in 1:5){
  des1[[i]] = rep(i,5*length(sivalst)*length(sivalsc)*length(yrvals)) #x5 = number of accuracy thresholds
}
des1=unlist(des1)

### reorder data for plotting and combine with data on which design is which

res3all3 = data.frame(cbind(c(res3all2[,c(1,4,3,2,5)]),rep(res3all2[,6],5),rep(res3all2[,7],5),rep(res3all2[,9],5),des1,rep(res3all2[,8],5)))
head(res3all3)
colnames(res3all3) = c("dev","sit","sic","acc","des1","yr")
res3all3$sit=as.numeric(res3all3$sit)
res3all3$sic=as.factor(res3all3$sic)
res3all3$acc=as.factor(res3all3$acc)
res3all3 = res3all3[rev(order(res3all3$sic)),]

### we decided to only plot the 10%, 30% and 50% accuracy thresholds and for no. time steps = 6
res3all4 = subset(res3all3, res3all3$acc!=0.2 & res3all3$acc!=0.4& res3all3$yr==6)


### use these to label facets
labels1 = c("�10%", "�30%","�50%")
labelsr = function(variable,value){return(labels1[value])}
library(ggplot2)
library(viridis)

### same graph as before but faceted by accuracy threshold
ggplot(res3all4, aes(x=sit,y=dev,group=interaction(des1,sic),fill=as.factor(des1))) +geom_line(col="black")+geom_point(aes(size=sic),pch=21,alpha=0.5) +
  scale_size_manual(name="Number of\ncontrol sites",values=c(2:8)) + scale_fill_manual(name=c("Design"),values=clrs,labels=c("RCT","BACI","CI","BA","After")) + facet_wrap(acc ~ ., nrow=1, labeller=labeller(.cols=labelsr)) +
  scale_x_continuous(name="Number of impact sites",limits=c(-2,52), breaks=c(1,5,10,25,50))+scale_y_continuous(name="Percentage of\nrepetitions with\ncorrect direction\n& within � x%\nof true effect", breaks=seq(0,100,10))+theme(axis.title.y  = element_text(angle=0, vjust=0.5, size=15),strip.text.x = element_text(size = 15),axis.text.x=element_text(size=15),axis.text.y=element_text(size=15),axis.title.x=element_text(size=15,hjust=0.5),legend.text = element_text(size=16),legend.title=element_text(size=16))+
  guides(fill = guide_legend(override.aes = list(size=10)))+
  theme(axis.title=element_text(size=16),aspect.ratio=3,strip.text = element_text(size=16),legend.text = element_text(size=15), legend.title = element_text(size=15))

##save plot if desired
#ggsave(filename = "results2.png",dpi = 300,width = 30,height = 30,units = "cm")

###suppl.info graph
ggplot(res3all3, aes(x=sit,y=dev,group=interaction(des1,sic),fill=as.factor(des1))) +geom_line(col="black")+geom_point(aes(size=sic),pch=21,alpha=0.8) +
  scale_size_manual(name="Number of\ncontrol sites",values=c(2:8)) + scale_fill_manual(name=c("Design"),values=clrs,labels=c("RCT","BACI","CI","BA","After")) + facet_wrap(acc + yr~ .,nrow=5) +
  scale_x_continuous(name="Number of impact sites",limits=c(-5,55), breaks=c(1,5,10,25,50))+scale_y_continuous(name="Percentage of\nrepetitions with\ncorrect direction\n& within � x%\nof true effect", breaks=seq(0,100,10))+theme(axis.title.y  = element_text(angle=0, vjust=0.5, size=15),strip.text.x = element_text(size = 13),axis.text.x=element_text(size=12),axis.text.y=element_text(size=12),axis.title.x=element_text(size=15,hjust=0.5),legend.text = element_text(size=16),legend.title=element_text(size=16))+
  guides(fill = guide_legend(override.aes = list(size=10)))+
  theme(axis.title=element_text(size=15),aspect.ratio=1,strip.text = element_text(size=15),legend.text = element_text(size=15), legend.title = element_text(size=14))
#ggsave(filename = "supplinfographsdirandacc.png",dpi = 300,width = 30,height = 30,units = "cm")


#### modelling results to produce equations of study design weights

### use results
res3all2 = data.frame(res3all2)
colnames(res3all2) = c(colnames(devall3)[1:5],"Sitest","Sitesc","yr","Accuracy")

###only for time step = 6
res3all4 = subset(res3all2,res3all2$yr==6)

### multiply percentages by 1000000 to find the number of simulation repetitions where effect size had correct sign and was within a given % of true effect size (perc success * 1000000 = no.successes)
### and number that didn't (100000000-number of successes = no.failures)
### 
res3all4[,1:5] = res3all4[,1:5]*1000000

###RCT ## consider all possible models
RCTm1 = glm(cbind(res3all4$RCT,100000000-res3all4$RCT)~log(res3all4$Sitest)+log(res3all4$Sitesc)+log(res3all4$Accuracy),  family='binomial')
RCTm2 = glm(cbind(res3all4$RCT,100000000-res3all4$RCT)~log(res3all4$Sitest)+log(res3all4$Sitesc)+(res3all4$Accuracy), family='binomial')
RCTm3 = glm(cbind(res3all4$RCT,100000000-res3all4$RCT)~log(res3all4$Sitest)+(res3all4$Sitesc)+(res3all4$Accuracy), family='binomial')
RCTm4 = glm(cbind(res3all4$RCT,100000000-res3all4$RCT)~(res3all4$Sitest)+(res3all4$Sitesc)+(res3all4$Accuracy), family='binomial')
RCTm5 = glm(cbind(res3all4$RCT,100000000-res3all4$RCT)~log(res3all4$Sitest)+(res3all4$Sitesc)+log(res3all4$Accuracy),  family='binomial')
RCTm6 = glm(cbind(res3all4$RCT,100000000-res3all4$RCT)~(res3all4$Sitest)+log(res3all4$Sitesc)+log(res3all4$Accuracy), family='binomial')
RCTm7 = glm(cbind(res3all4$RCT,100000000-res3all4$RCT)~(res3all4$Sitest)+log(res3all4$Sitesc)+(res3all4$Accuracy), family='binomial')
RCTm8 = glm(cbind(res3all4$RCT,100000000-res3all4$RCT)~(res3all4$Sitest)+(res3all4$Sitesc)+log(res3all4$Accuracy), family='binomial')

###check AICs
AIC(RCTm1,RCTm2,RCTm3,RCTm4,RCTm5,RCTm6,RCTm7,RCTm8)

###best model
summary(RCTm1)

##units are log odds to covert to probabilities you need to use below function or equation
predict(RCTm1, newdata=res3all4,type="response")
#1/(1+exp(-(1.730e+00 + 2.973e-02*log(nI) + 2.868e-02*log(nC) + 1.309e+00*log(A))))


###find pseudo-R2 value using null model
RCTnull = glm(cbind(res3all4$RCT,100000000-res3all4$RCT)~1, family='binomial')
1 - logLik(RCTm1)/logLik(RCTnull)


#### Repeat for all the other designs
####BACI 
BACIm1 = glm(cbind(res3all4$BACI,100000000-res3all4$BACI)~log(res3all4$Sitest)+log(res3all4$Sitesc)+log(res3all4$Accuracy), family='binomial')
BACIm2 = glm(cbind(res3all4$BACI,100000000-res3all4$BACI)~log(res3all4$Sitest)+log(res3all4$Sitesc)+(res3all4$Accuracy), family='binomial')
BACIm3 = glm(cbind(res3all4$BACI,100000000-res3all4$BACI)~log(res3all4$Sitest)+(res3all4$Sitesc)+(res3all4$Accuracy), family='binomial')
BACIm4 = glm(cbind(res3all4$BACI,100000000-res3all4$BACI)~(res3all4$Sitest)+(res3all4$Sitesc)+(res3all4$Accuracy), family='binomial')
BACIm5 = glm(cbind(res3all4$BACI,100000000-res3all4$BACI)~log(res3all4$Sitest)+(res3all4$Sitesc)+log(res3all4$Accuracy),  family='binomial')
BACIm6 = glm(cbind(res3all4$BACI,100000000-res3all4$BACI)~(res3all4$Sitest)+log(res3all4$Sitesc)+log(res3all4$Accuracy), family='binomial')
BACIm7 = glm(cbind(res3all4$BACI,100000000-res3all4$BACI)~(res3all4$Sitest)+log(res3all4$Sitesc)+(res3all4$Accuracy), family='binomial')
BACIm8 = glm(cbind(res3all4$BACI,100000000-res3all4$BACI)~(res3all4$Sitest)+(res3all4$Sitesc)+log(res3all4$Accuracy), family='binomial')

AIC(BACIm1,BACIm2,BACIm3,BACIm4,BACIm5,BACIm6,BACIm7,BACIm8)

summary(BACIm1)


BACInull = glm(cbind(res3all4$BACI,100000000-res3all4$BACI)~1, family='binomial')
1 - logLik(BACIm1)/logLik(BACInull)


####BA
BAm1 = glm(cbind(res3all4$BA,100000000-res3all4$BA)~log(res3all4$Sitest)+log(res3all4$Accuracy), family='binomial')
BAm2 = glm(cbind(res3all4$BA,100000000-res3all4$BA)~log(res3all4$Sitest)+(res3all4$Accuracy), family='binomial')
BAm3 = glm(cbind(res3all4$BA,100000000-res3all4$BA)~(res3all4$Sitest)+log(res3all4$Accuracy), family='binomial')
BAm4 = glm(cbind(res3all4$BA,100000000-res3all4$BA)~(res3all4$Accuracy), family='binomial')
BAm5 = glm(cbind(res3all4$BA,100000000-res3all4$BA)~(res3all4$Sitest)+(res3all4$Accuracy), family='binomial')
BAm6 = glm(cbind(res3all4$BA,100000000-res3all4$BA)~log(res3all4$Accuracy), family='binomial')
BAm7 = glm(cbind(res3all4$BA,100000000-res3all4$BA)~log(res3all4$Sitest), family='binomial')
BAm8 = glm(cbind(res3all4$BA,100000000-res3all4$BA)~(res3all4$Sitest), family='binomial')

AIC(BAm1,BAm2,BAm3,BAm4,BAm5,BAm6,BAm7,BAm8)

summary(BAm1)

BAnull = glm(cbind(res3all4$BA,100000000-res3all4$BA)~1, family='binomial')
1 - logLik(BAm1)/logLik(BAnull)


####CI
CIm1 = glm(cbind(res3all4$CI,100000000-res3all4$CI)~log(res3all4$Sitest)+log(res3all4$Sitesc)+log(res3all4$Accuracy), family='binomial')
CIm2 = glm(cbind(res3all4$CI,100000000-res3all4$CI)~log(res3all4$Sitest)+log(res3all4$Sitesc)+(res3all4$Accuracy), family='binomial')
CIm3 = glm(cbind(res3all4$CI,100000000-res3all4$CI)~log(res3all4$Sitest)+(res3all4$Sitesc)+(res3all4$Accuracy), family='binomial')
CIm4 = glm(cbind(res3all4$CI,100000000-res3all4$CI)~(res3all4$Sitest)+log(res3all4$Sitesc)+log(res3all4$Accuracy), family='binomial')
CIm5 = glm(cbind(res3all4$CI,100000000-res3all4$CI)~(res3all4$Sitest)+log(res3all4$Sitesc)+(res3all4$Accuracy), family='binomial')

CIm6 = glm(cbind(res3all4$CI,100000000-res3all4$CI)~(res3all4$Sitest)+(res3all4$Sitesc)+log(res3all4$Accuracy), family='binomial')
CIm7 = glm(cbind(res3all4$CI,100000000-res3all4$CI)~log(res3all4$Sitest)+(res3all4$Sitesc)+log(res3all4$Accuracy), family='binomial')
CIm8 = glm(cbind(res3all4$CI,100000000-res3all4$CI)~(res3all4$Sitest)+(res3all4$Sitesc)+(res3all4$Accuracy), family='binomial')
CIm9 = glm(cbind(res3all4$CI,100000000-res3all4$CI)~log(res3all4$Sitesc)+log(res3all4$Accuracy), family='binomial')
CIm10 = glm(cbind(res3all4$CI,100000000-res3all4$CI)~log(res3all4$Sitest)+log(res3all4$Accuracy), family='binomial')
CIm11 = glm(cbind(res3all4$CI,100000000-res3all4$CI)~log(res3all4$Sitesc)+log(res3all4$Sitest), family='binomial')

CIm12 = glm(cbind(res3all4$CI,100000000-res3all4$CI)~(res3all4$Sitesc)+log(res3all4$Accuracy), family='binomial')
CIm13 = glm(cbind(res3all4$CI,100000000-res3all4$CI)~(res3all4$Sitest)+log(res3all4$Accuracy), family='binomial')
CIm14 = glm(cbind(res3all4$CI,100000000-res3all4$CI)~(res3all4$Sitesc)+log(res3all4$Sitest), family='binomial')

CIm15 = glm(cbind(res3all4$CI,100000000-res3all4$CI)~log(res3all4$Sitesc)+(res3all4$Accuracy), family='binomial')
CIm16 = glm(cbind(res3all4$CI,100000000-res3all4$CI)~log(res3all4$Sitest)+(res3all4$Accuracy), family='binomial')
CIm17 = glm(cbind(res3all4$CI,100000000-res3all4$CI)~log(res3all4$Sitesc)+(res3all4$Sitest), family='binomial')

CIm18 = glm(cbind(res3all4$CI,100000000-res3all4$CI)~(res3all4$Sitesc)+(res3all4$Accuracy), family='binomial')
CIm19 = glm(cbind(res3all4$CI,100000000-res3all4$CI)~(res3all4$Sitest)+(res3all4$Accuracy), family='binomial')
CIm20 = glm(cbind(res3all4$CI,100000000-res3all4$CI)~(res3all4$Sitesc)+(res3all4$Sitest), family='binomial')

CIm21 = glm(cbind(res3all4$CI,100000000-res3all4$CI)~(res3all4$Sitesc), family='binomial')
CIm22 = glm(cbind(res3all4$CI,100000000-res3all4$CI)~(res3all4$Sitest), family='binomial')
CIm23 = glm(cbind(res3all4$CI,100000000-res3all4$CI)~(res3all4$Accuracy), family='binomial')

CIm24 = glm(cbind(res3all4$CI,100000000-res3all4$CI)~log(res3all4$Sitesc), family='binomial')
CIm25 = glm(cbind(res3all4$CI,100000000-res3all4$CI)~log(res3all4$Sitest), family='binomial')
CIm26 = glm(cbind(res3all4$CI,100000000-res3all4$CI)~log(res3all4$Accuracy), family='binomial')

AIC(CIm1,CIm2,CIm3,CIm4,CIm5,CIm6,CIm7,CIm8,CIm9,CIm10,
    CIm11,CIm12,CIm13,CIm14,CIm15,CIm16,CIm17,CIm18,CIm19,CIm20,
    CIm21,CIm22,CIm23,CIm24,CIm25,CIm26)
summary(CIm1)

CInull = glm(cbind(res3all4$CI,100000000-res3all4$CI)~1, family='binomial')
1 - logLik(CIm1)/logLik(CInull)


####AFTER
Afterm1 = glm(cbind(res3all4$After,100000000-res3all4$After)~log(res3all4$Sitest)+log(res3all4$Accuracy), family='binomial')
Afterm2 = glm(cbind(res3all4$After,100000000-res3all4$After)~log(res3all4$Sitest)+(res3all4$Accuracy), family='binomial')
Afterm3 = glm(cbind(res3all4$After,100000000-res3all4$After)~(res3all4$Sitest)+log(res3all4$Accuracy), family='binomial')
Afterm4 = glm(cbind(res3all4$After,100000000-res3all4$After)~(res3all4$Accuracy), family='binomial')
Afterm5 = glm(cbind(res3all4$After,100000000-res3all4$After)~(res3all4$Sitest)+(res3all4$Accuracy), family='binomial')
Afterm6 = glm(cbind(res3all4$After,100000000-res3all4$After)~log(res3all4$Accuracy), family='binomial')
Afterm7 = glm(cbind(res3all4$After,100000000-res3all4$After)~log(res3all4$Sitest), family='binomial')
Afterm8 = glm(cbind(res3all4$After,100000000-res3all4$After)~(res3all4$Sitest), family='binomial')

AIC(Afterm1,Afterm2,Afterm3,Afterm4,Afterm5,Afterm6,Afterm7,Afterm8)

summary(Afterm1)

Afternull = glm(cbind(res3all4$After,100000000-res3all4$After)~1, family='binomial')
1 - logLik(Afterm1)/logLik(Afternull)
```
