## Supporting Information for "Simple study designs in ecology produce inaccurate estimates of biodiversity responses"

### Supporting Information Section A: Empirical parameterisation

The major parameters in this simulation include: the change between the before and after periods in both impact sites ($I$) and control sites ($C$) and the difference between impact and control group means before intervention ($d_{CIB}$). The values of these three parameters were taken from 47 BACI datasets (Christie et al., unpub) whereby the proportional change or difference (Table 3 main text) was extracted for 2002 effect sizes. We collated 45 datasets using a full search of the Web of Science with the search terms: [‘BACI’] OR [‘Before-After Control-Impact’] on the 18th December 2017. The search returned 674 results and we then refined this by selecting only ‘Article’ as a document type and only the following Web of Science Categories: ‘Ecology’, ‘Marine Freshwater Biology’, ‘Biodiversity Conservation’, ‘Fisheries’, ‘Oceanography’, ‘Forestry’, ‘Zoology’, Ornithology’, ‘Biology’, ‘Plant Sciences’, ‘Entomology’, ‘Remote Sensing’, ‘ Toxicology’ and ‘Soil Science’. We were then left with 579 results. To be realistic about obtaining the data, we restricted the year of publication to 2002 (15 years prior to search), which reduced the number to 542. We then read the abstracts of all papers that we could access and excluded any studies that did not test the effect of an ecological intervention or threat using a BACI design with abundance, cover or density metrics of any taxon. This left 96 studies for which we then contacted the corresponding authors to ask for their raw data and received 45 datasets. We also collected 2 datasets from contacts and collaborators. We found that 25 datasets tested the effects of threats and 22 tested the effects of interventions.

It is possible that the datasets that we used were subject to publication bias as studies that show statistically significant positive effects are more likely to be published (Chalmers 1990; Easterbrook et al. 1991). In this case, a BACI dataset is more likely to give a significant result if there are large, significant opposing changes in impact and control sites before versus after the impact (a time-impact interaction). Therefore, we might expect more BACI datasets to be published that show large, possibly unrepresentative changes in impacts and/or controls. To limit this bias, we tried to extract all the raw data from datasets, rather than just the data that was the major focus of the publication. We also tried to counteract any extreme values from datasets by only considering data within the Interquartile Range (IQR) of each parameter, separately. These 25% and 75% quantiles then became the minimum and maximum values of the subsetted data (number of values per parameter: $I$=1320; $C$=1206; $d_{CIB}$=1166). We randomly sampled this subsetted data for each parameter 1000 times with replacement (Table 3 main text) and used these values to create 1000 unique simulation scenarios.

### Supporting Tables 1-5

Table 1 - Models considered in model selection process for finding weighting equation for RCT design.$N_I$= Number of Impact sites, $N_C$= Number of Control sites, $A$= Accuracy threshold (e.g. $\pm 10\%$= 0.1).

| | Parameters | AIC |
|---|---|---|
| Best model | $Intercept + ln(N_I) + ln(N_C) + ln(A)$ | 1995902 |
| Model A | $Intercept + ln(N_I) + ln(N_C) + (A)$ | 77379448 |
| Model B | $Intercept + ln(N_I) + (N_C) + ln(A)$ | 4231749 |
| Model C | $Intercept + (N_I) + ln(N_C) + ln(A)$ | 4390830 |
| Model D | $Intercept + ln(N_I) + (N_C) + (A)$ | 79602006 |
| Model E | $Intercept + (N_I) + (N_C) + ln(A)$ | 6626242 |
| Model F | $Intercept + (N_I) + ln(N_C) + (A)$ | 79760180 |
| Model G | $Intercept + (N_I) + (N_C) + (A)$ | 81982310 |

Table 2 - Models considered in model selection process for finding weighting equation for BACI design.

| | Parameters | AIC |
|---|---|---|
| Best model | $Intercept + ln(N_I) + ln(N_C) + ln(A)$ | 68044756 |
| Model A | $Intercept + ln(N_I) + ln(N_C) + (A)$ | 106675117 |
| Model B | $Intercept + ln(N_I) + (N_C) + ln(A)$ | 154570805 |
| Model C | $Intercept + (N_I) + ln(N_C) + ln(A)$ | 190252540 |
| Model D | $Intercept + ln(N_I) + (N_C) + (A)$ | 192738538 |
| Model E | $Intercept + (N_I) + (N_C) + ln(A)$ | 275789163 |
| Model F | $Intercept + (N_I) + ln(N_C) + (A)$ | 228246468 |
| Model G | $Intercept + (N_I) + (N_C) + (A)$ | 313321891 |

Table 3 - Models considered in model selection process for finding weighting equation for BA design.

| | Parameters | AIC |
|---|---|---|
| Best model | $Intercept + ln(N_I) + ln(A)$ | 126468.1 |
| Model A | $Intercept + ln(N_I) + (A)$ | 32655553.6 |
| Model B | $Intercept + (N_I) + ln(A)$ | 334893.0 |
| Model C | $Intercept + (N_I) + (A)$ | 32864140.3 |
| Model D | $Intercept + ln(N_I)$ | 655710676.4 |
| Model E | $Intercept + (N_I)$ | 655908966.3 |
| Model F | $Intercept + ln(A)$ | 510145.2 |
| Model G | $Intercept + (A)$ | 33039522.2 |

Table 4 - Models considered in model selection process for finding weighting equation for CI design.

| | Parameters | AIC |
|---|---|---|
| Best model | $Intercept + ln(N_I) + ln(N_C) + ln(A)$ | 270896.2 |
| Model A | $Intercept + ln(N_I) + ln(N_C) + (A)$ | 19981372.1 |
| Model B | $Intercept + ln(N_I) + (N_C) + ln(A)$ | 276828.6 |
| Model C | $Intercept + (N_I) + ln(N_C) + ln(A)$ | 276681.0 |
| Model D | $Intercept + ln(N_I) + (N_C) + (A)$ | 19987312.1 |
| Model E | $Intercept + (N_I) + (N_C) + ln(A)$ | 282613.4 |
| Model F | $Intercept + (N_I) + ln(N_C) + (A)$ | 19987164.3 |
| Model G | $Intercept + (N_I) + (N_C) + (A)$ | 19993104.4 |
| Model H | $Intercept + ln(N_I) + ln(N_C)$ | 503366818.6 |
| Model I | $Intercept + ln(N_I) + ln(A)$ | 281386.8 |
| Model J | $Intercept + ln(N_C) + ln(A)$ | 280817.8 |
| Model K | $Intercept + (N_I) + ln(N_C)$ | 503372387.2 |

| | | |
|---|---|---|
| Model L | $Intercept\ + (N_I)\ +\ ln(A)$ | 287171.6 |
| Model M | $Intercept\ + (N_C)\ + ln(A)$ | 286750.2 |
| Model N | $Intercept\ + ln(N_I)\ +\ (N_C)$ | 503372529.4 |
| Model O | $Intercept\ + ln(N_I)\ + (A)$ | 19991876.2 |
| Model P | $Intercept\ + ln(N_C)\ + (A)$ | 19991306.4 |
| Model Q | $Intercept\ + (N_I)\ +\ (N_C)$ | 503378098.0 |
| Model R | $Intercept\ + (N_I)\ + (A)$ | 19997668.4 |
| Model S | $Intercept\ + (N_C)\ + (A)$ | 19997246.5 |
| Model T | $Intercept\ + ln(N_C)$ | 503376369.2 |
| Model U | $Intercept\ + ln(N_I)$ | 503376917.0 |
| Model V | $Intercept\ + ln(A)$ | 291308.4 |
| Model W | $Intercept\ + (N_C)$ | 503382080.0 |
| Model X | $Intercept\ + (N_I)$ | 503382485.6 |
| Model Y | $Intercept\ + (A)$ | 20001810.5 |

Table 5 - Models considered in model selection process for finding weighting equation for After design.

| | Parameters | AIC |
|---|---|---|
| Best model | $Intercept\ +\ ln(N_I)\ +\ ln(A)$ | 1103236 |
| Model A | $Intercept\ +\ ln(N_I)\ +\ (A)$ | 9143502 |
| Model B | $Intercept\ +\ (N_I)\ +\ ln(A)$ | 1468954 |
| Model C | $Intercept\ + (N_I)\ +\ (A)$ | 9509817 |
| Model D | $Intercept\ + ln(N_I)$ | 330771704 |
| Model E | $Intercept\ + (N_I)$ | 331128413 |
| Model F | $Intercept\ + ln(A)$ | 1841665 |
| Model G | $Intercept\ +\ (A)$ | 9883134 |

### Supporting Figure 1

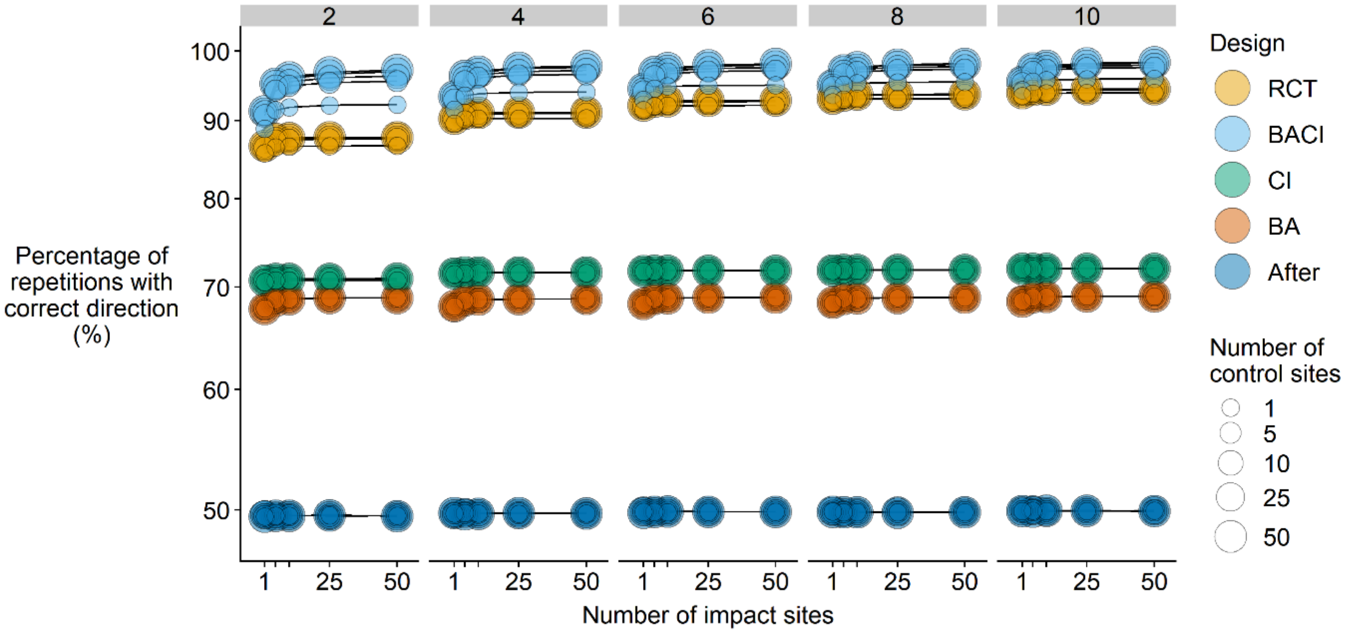

Fig.1 - Accuracy of different designs measured by the percentage of simulation repetitions in which the design's effect size had the correct direction for multiple numbers of time steps simulated (T = 2, 4, 6, 8 or 10) and levels of spatial replication (control and impact sites separately). Y-axis uses a $\log_{10}$ scale and circle size denotes the number of control sites. See Table 1 in main text for the definition of each design.

### Supporting Figure 2

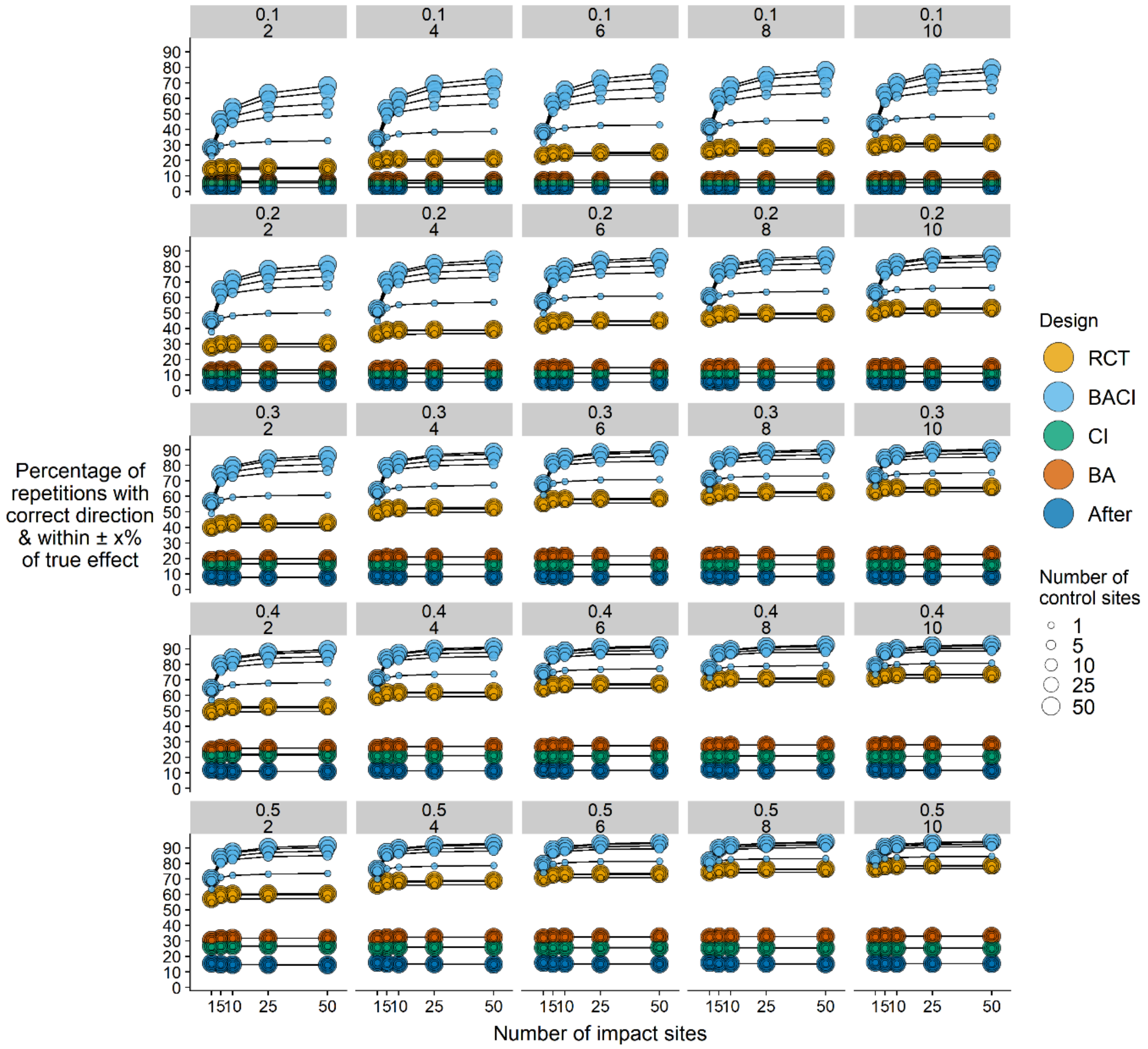

Fig.2 - Accuracy of different designs measured by the percentage of simulation repetitions in which the design's effect size was both within 10, 30 or 50% of the true effect size and had the correct direction for multiple levels of spatial replication (control and impact sites separately). Graphs show all possible combinations of accuracy thresholds (0.1 = ±10%, 0.3 = ±30% or 0.5 = ±50%) and time steps simulated (*T* = 2, 4, 6, 8 or 10). Circle size denotes the number of control sites. See Table 1 in main text for the definition of each design.

### Description of Supporting Code zip file

The Supporting code is an R script file containing all the code necessary to run the simulation using imported empirical estimates of *I*, *C* and $d_{CIB}$ if formatted as found in the *empiricaldata.csv* file. Details on how empirical estimates were calculated can be found in Table 2 of the main text. The columns in the csv file are as follows:

- DatasetID – Anonymised ID of individual raw datasets from which the parameter estimate data was calculated (note there are more than 47 IDs as some datasets had multiple components).

- EFID – Anonymised ID of individual data source within raw datasets – for example, we used raw data for multiple species from each dataset to calculate parameter estimates.

- Type – Type of impact the dataset considered which is either an ‘Intervention’ or a ‘Threat’.

- Cht – Change in impact sites between before and after periods ($p_I$ in main text).

- Chc - Change in control between before and after periods ($p_C$ in main text).

- tcd - Average value of control sites as a proportion of the average value of impact sites in the before period ($p_{CIB}$ in main text).

The R script also contains code for reproducing our figures and for generating the accuracy weight equations in the main text.
